# Interaction of yeast V-ATPase with TLDc protein Rtc5p

**DOI:** 10.1101/2025.05.24.655954

**Authors:** Md. Murad Khan, Roshanak Ebrahimi, Rebecca A. Oot, Stephan Wilkens

**Affiliations:** Department of Biochemistry and Molecular Biology, SUNY Upstate Medical University, Syracuse, NY 13210, USA

**Keywords:** Vacuolar H^+^-ATPase, Rtc5p, TLDc domain, Reversible disassembly, Protein structure

## Abstract

The eukaryotic vacuolar H^+^-ATPase (V-ATPase) is regulated by reversible disassembly into autoinhibited V_1_-ATPase and V_o_ proton channel subcomplexes, a mode of regulation conserved from yeast to humans. While signals that govern V-ATPase assembly have been studied in the cellular context, the molecular mechanisms of the process at the level of the enzyme remain poorly understood. We recently discovered that Oxr1p, one of the two TLDc domain proteins in yeast, is essential for rapid V-ATPase *disassembly in vivo*. How the second TLDc protein, Rtc5p, functions in reversible disassembly, however, is less clear. Here we find that Rtc5p promotes assembly of functional holo V-ATPase from purified V_1_ and V_o_ subcomplexes *in vitro*. CryoEM structures show that Rtc5p’s TLDc domain binds the C-terminal domain of the V_1_-B subunit, with Rtc5p’s C-terminal a-helix inserting into the catalytic hexamer, thereby opening a second catalytic site distal to its binding site. Unlike Oxr1p, however, which when deleted produces a distinct phenotype, Rtc5p does not appear to be essential for glucose driven enzyme (re)assembly, hinting at the presence of multiple assembly pathways *in vivo*.

## Introduction

Vacuolar ATPases (V-ATPases; V_1_V_o_-ATPases) are ubiquitous multi-subunit proton pumps responsible for acidifying organelles in virtually all eukaryotic cells^1–4^. The differential pH established by the V-ATPase governs organellar identity and drives fundamental cellular processes, including pH and ion homeostasis, lysosomal degradation, hormone secretion^5^, and neurotransmitter release^6^. V-ATPase is also found at the plasma membrane in specialized tissues, such as osteoclasts and renal α-intercalated cells, where movement of protons across the plasma membrane is critical for bone resorption and acid removal, respectively. In addition, V-ATPase’s proton pumping function plays a critical role in vital signaling pathways such as notch^7^, mTOR^8^ and Wnt^9^. V-ATPase is highly conserved from yeast to humans, making yeast a powerful model system to study the enzyme’s structure and mechanism. V-ATPase is assembled from two subcomplexes, a peripheral ATPase (V_1_) and a membrane embedded proton channel (V_o_)^10–12^. ATP hydrolysis on V_1_ is coupled to transmembrane proton flow across V_o_. The V_1_ is composed of eight different subunits A-H in varying stoichiometries (A_3_B_3_CDE_3_FG_3_H), with three catalytic nucleotide binding sites located at subunit AB interfaces. The V_o_ contains seven different polypeptides in a ratio of *ac*_8_*c*′*c*′′*def* (for the yeast enzyme; *ac*_9_*c*′′*def* in mammalian V-ATPase). The integral *c* subunits (“proteolipids”, *c*, *c*′, *c*′′) are arranged in a ten membered ring (*c*-ring) that shares an interface with the membrane embedded C-terminal domain of subunit *a* (*a*_CT_). Sequential binding and hydrolysis of ATP in the three catalytic sites of V_1_ is coupled to rotation of a central rotor subcomplex composed of V_1_ subunits DF, V_o_ subunit *d*, and the *c*-ring. Each proteolipid chain contains a conserved glutamate residue exposed to the lipid bilayer that carries protons between cytosolic and luminal aqueous half channels located in *a*_CT_. Three peripheral stalks composed of subunit EG heterodimers (EG1, EG2 and EG3), together with the collar subunits H and C and the N-terminal domain of subunit *a* (*a*_NT_) keep the V_1_ fixed relative to V_o_ for efficient energy coupling. V_1_ is a stepping motor: each ATP binding/hydrolysis event drives the central rotor in steps of 120°, with each of the three steps coupled to the transfer of ∼3.3 protons across the *c*-ring:*a*_CT_ interface^13^. The rotary states of the V-ATPase have been visualized using cryo electron microscopy (cryoEM) of the enzymes from yeast^14,15^ and higher organisms^16–18^.

V-ATPase activity is tightly regulated, most prominently by a mechanism referred to as *reversible disassembly*. In yeast, for example, the majority of V-ATPases are assembled and active in the presence of glucose. Upon glucose withdrawal, however, subunit C is rapidly released into the cytosol, with subsequent separation and inactivation of V_1_ and V_o_ subcomplexes (**Fig. 1A**)^19–21^. When glucose levels are restored, subunit C and V_1_ and V_o_ reassociate to form active holoenzymes. Studies in yeast have shown that efficient *disassembly* requires catalytically active enzymes and intact microtubules^22,23^. More recently, we found that the *in vivo* disassembly process requires the presence of *Oxidation Resistance 1 protein* (Oxr1p), and from *in vitro* experiments, we showed that Oxr1p binding to the holoenzyme is sufficient to cause disassembly^15,24,25^. On the other hand, efficient (*re*)*assembly* depends on the heterotrimeric chaperone complex called *RAVE* (**<u>R</u>**egulator of the H^+^-**<u>A</u>**TPase of **<u>V</u>**acuolar and **<u>E</u>**ndosomal membranes), which delivers V_1_ and subunit C back to the vacuolar membrane in a glucose dependent manner^26–28^. RAVE, however, is not sufficient for facilitating enzyme assembly from purified components *in vitro*^29^, indicating that additional factors are required for the process.

**Figure 1.**
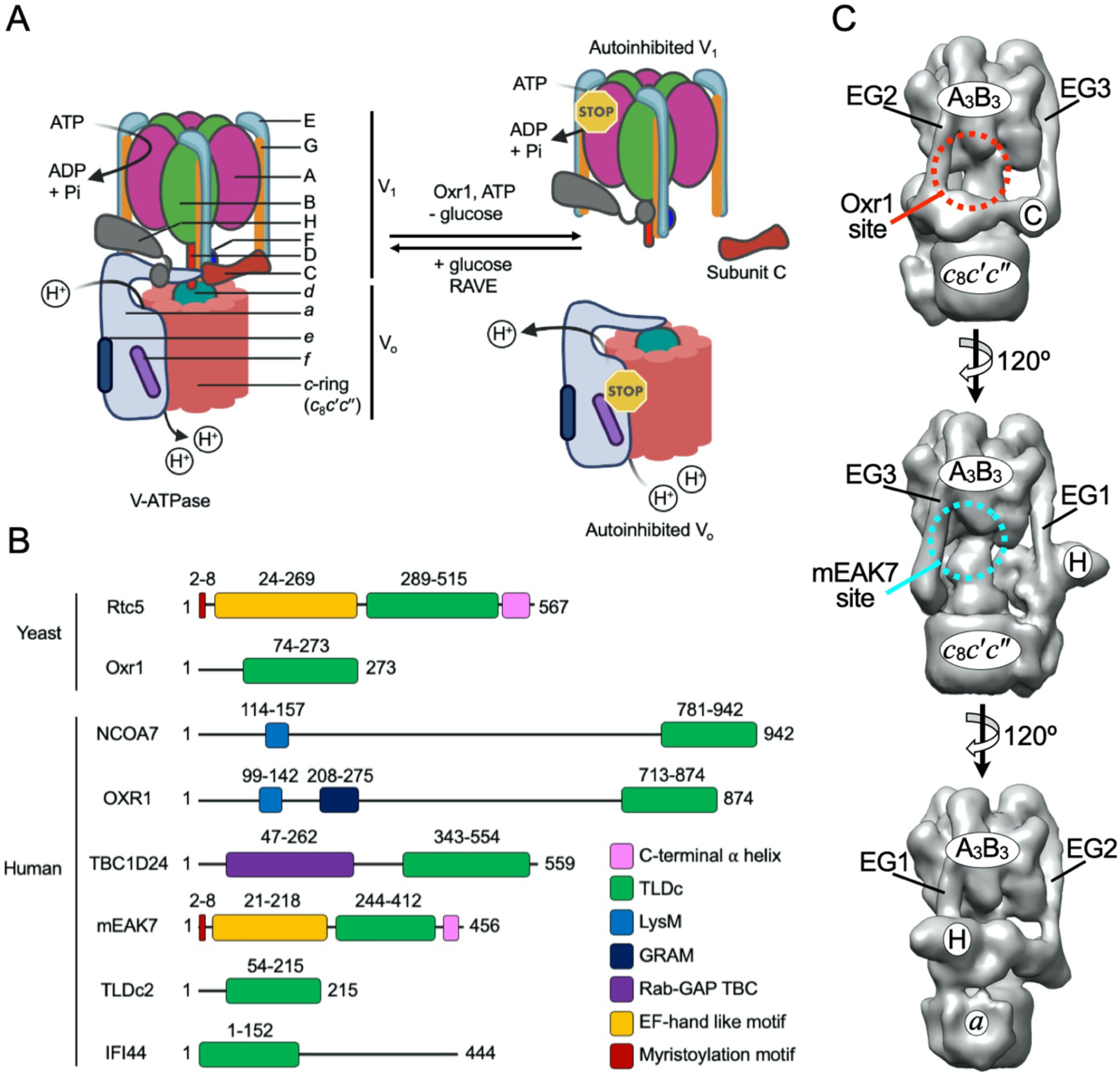
Regulation of V-ATPase activity by reversible disassembly and domain architecture of TLDc proteins. **A.** Schematic of V-ATPase regulation by reversible disassembly. In yeast, under glucose starvation, TLDc protein Oxr1p disassembles the enzyme into V_1_ and V_o_ subcomplexes, with both V_1_ and V_o_ becoming autoinhibited. Restoration of glucose leads to reassembly of functional holoenzymes. Efficient reassembly requires an assembly chaperone called the *RAVE complex*. **B.** Domain architecture of TLDc domain proteins from yeast and humans. All members of the family have a C-terminal TLDc domain except IFI44, where the TLDc domain is near the N-terminus. Domain boundaries are from UniProt^36^: Yeast Rtc5p (Q12108) and Oxr1p (Q08952); human mEAK7 (Q6P9B6), OXR1 (Q8N573), NCOA7 (Q8NI08), TBC1D24 (Q9ULP9), TLDC2 (A0PJX2) and IFI44 (Q8TCB0). Myristoylation and EF-hand motifs are from Johnson et al. (1994)^37^ and Tan et al. (2022)^38^, respectively. **C.** Views of a V-ATPase surface representation highlighting the binding sites for Oxr1p (top panel, red dashed circle) and mEAK7 (middle panel, cyan dashed circle). Oxr1p binds the B subunit C-terminal domain next to peripheral stator EG2 and subunit C, and mEAK7 binds the B subunit next to EG3. TLDc proteins have not yet been seen to bind the B subunit next to EG1 (bottom panel). The surface representation was generated using the coordinate file for the yeast V-ATPase in rotary state 1 (PDBID: 7FDA^15^).

Yeast Oxr1p belongs to the family of *TLDc* (Tre2/Bub2/Cdc16 (**<u>T</u>**BC), lysin motif (**<u>L</u>**ysM), **<u>D</u>**omain **<u>c</u>**atalytic) domain proteins^30^ that are conserved from yeast to humans. The family is characterized by the presence of a C-terminal TLDc domain, and while some consist entirely of the TLDc motif, others contain additional N-terminal domains (**Fig. 1B**). So far, two TLDc proteins have been identified in yeast, Oxr1p^31^ and *Restriction of Telomere Capping 5 protein* (Rtc5p)^32,33^, whereas at least six have been described in higher organisms, including humans (NCOA7, OXR1, TLDC2, TBC1D24, mEAK7, and IFI44)^30^. TLDc proteins have been recognized as V-ATPase binding proteins^34^, and we recently showed that four of the human TLDc proteins (NCOA7, OXR1, TLDC2 and TBC1D24) disassemble purified human V-ATPase in an ATP dependent manner, but that a fifth one, mEAK7, activates the enzyme^35^.

CryoEM structural analysis of Oxr1p and mEAK7 bound V-ATPase complexes revealed that the two TLDc proteins bind the enzyme at different but equivalent sites in a rotary state-dependent manner^15,38,39^. Whereas Oxr1p binds the C-terminal domain of the B subunit next to peripheral stator EG2 in rotary state 1, mEAK7 interacts with the B subunit next to EG3 in rotary state 2, with the two sites hereafter referred to as *Oxr1 site* and *mEAK7 site*, respectively (**Fig. 1C**).

Yeast Rtc5p is a structural homolog of mEAK7, both have N-terminal myristoylation and EF-hand-like motifs followed by a TLDc domain and a C-terminal a helical segment^33^ (**Fig. 1B**). Yeast Rtc5p has recently been proposed to act as yet another disassembly factor, although it was suggested to be less efficient as compared to Oxr1p^33^. Here we investigate Rtc5p’s molecular role in regulating the assembly state of the yeast V-ATPase. We show that the interaction between V-ATPase and recombinant Rtc5p does not lead to enzyme inhibition or disassembly *in vitro*. CryoEM analysis reveals that Rtc5p can bind intact V-ATPase both in the Oxr1p binding site and the mEAK7 site in a rotary state dependent manner, and that the interaction is disrupted in the presence of nucleotides. We further find that while Rtc5p binding to wild type V_1_ does not relieve autoinhibition, it primes the V_1_ for assembly with both purified V_o_, and V_o_ in vacuolar membranes. *In vivo* studies, however, show that glucose driven V-ATPase (re)assembly can occur in the absence of Rtc5p, indicating involvement of additional factors and/or alternative pathways.

## Results

### Rtc5p has no apparent effect on holoenzyme activity *in vitro*

We previously reported that Oxr1p, one of the two TLDc domain proteins in yeast, catalyzes the disassembly step of V-ATPase regulation by reversible disassembly^15,24^. A recent study by Klössel et al. concluded that the other yeast TLDc protein, Rtc5p, also contributes to V-ATPase disassembly^33^. The molecular details of Rtc5p’s proposed role in the process, however, are not yet known. To determine if Rtc5p affected V-ATPase activity or assembly directly, we expressed the protein in *E. coli* for biochemical and biophysical experiments (**Appendix Fig. S1A**). Size exclusion chromatography (SEC) showed that Rtc5p elutes at its expected mass of ∼65 kDa, indicating that the protein is monomeric in solution (**Appendix Fig. S1B**). We then incubated purified vacuoles with or without Rtc5p at a 1:5 w/w ratio over vacuolar protein for 30 min at room temperature followed by ATPase activity assays and found that under these conditions, Rtc5p has no significant effect on the enzyme’s activity (**Fig 2A**). Previously, we have shown that Oxr1p mediated V-ATPase disassembly is a slow process unless ATP is added^24^. We therefore repeated the experiment in the presence of ATP, and while Oxr1p leads to complete inhibition under this condition, we observed only slightly less activity of vacuoles after incubation with ATP, irrespective of whether Rtc5p was present or not **(Fig 2A**). To test whether the slight inhibition with Rtc5p and ATP is due to enzyme disassembly, we incubated purified vacuoles with or without Rtc5p (at the same 1:5 ratio) and ATP for 30 min at room temperature before pelleting vacuoles and probing supernatants for the presence of V_1_ subunits using western blotting. The analysis showed only trace amounts of V_1_ subunit A in the supernatants of the vacuoles treated with ATP, regardless of whether Rtc5p was present or not (**Fig 2B**). This indicated that Rtc5p did not cause enzyme disassembly under these conditions. Conducting the experiment with Oxr1p + ATP, however, led to almost complete release of V_1_ from vacuoles as expected based on our previous work^24^. To summarize, Rtc5p, in contrast to Oxr1p, does not cause inhibition or disassembly of the V-ATPase on vacuolar membranes under the conditions tested.

**Figure 2.**
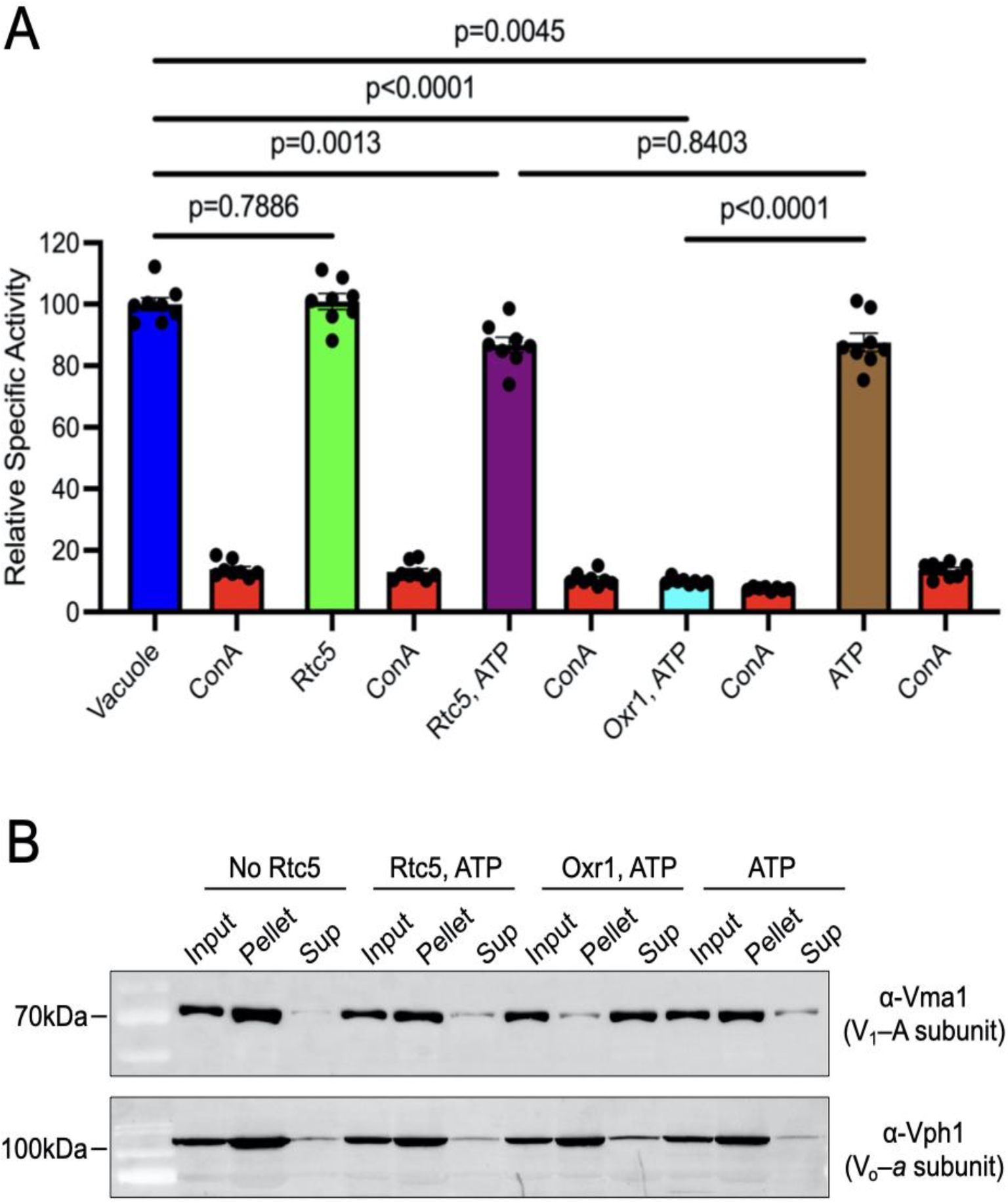
Rtc5p has no apparent effect on activity and assembly state of V-ATPase. **A.** Purified wild type vacuoles were incubated with an estimated 1:5 ratio of Rtc5p over vacuolar protein (w/w) in absence (green) and presence (purple) of 4mM MgATP for 30 min at room temperature followed by measuring ATPase activity. Relative activities were normalized against the starting activity (blue). As a positive control of activity inhibition, vacuoles were incubated with Oxr1p (at a ratio of 1:10 (w/w) to vacuoles) in presence of MgATP (cyan). The effect of MgATP alone was also determined (brown). The V-ATPase specific inhibitor Concanamycin A (ConA) was included as an additional control (red). Individual data points of eight measurements from two to three biological preparations are shown. Data are presented as mean ± SEM. **B.** Purified vacuoles were incubated for 30 min at room temperature under the indicated conditions. Rtc5p and Oxr1p were added at ratios of 1:5 and 1:10 (w/w) to vacuoles, respectively, and MgATP was added to a final concentration of 4 mM. Vacuolar membranes were pelleted by ultracentrifugation, and supernatant (*Sup*) and pellet (*Pellet*) samples were analyzed by western blotting using an α-Vma1 (V_1_ subunit A) antibody. Probing with an α-Vph1 (V_o_ subunit *a*) antibody was included as a control. A representative blot from three independent experiments is shown.

### Rtc5p is not required for glucose starvation-induced V-ATPase disassembly *in vivo*

To investigate Rtc5p’s putative role in V-ATPase disassembly *in vivo*, we deleted the *RTC5* locus to generate the *rtc5Δ* strain. In yeast, V-ATPase loss of function produces a conditional lethal phenotype (Vma^−^)^40^ characterized by a failure to grow at alkaline pH in presence of Ca^2+^. Consistent with the study by Klössel et al.^33^, we observed that deletion of *RTC5* did not produce a Vma^−^ growth phenotype (**Appendix Figure 2**). We then used fluorescence microscopy to monitor the assembly state of the V-ATPase upon glucose deprivation in yeast cells containing C-terminally mNeonGreen (mNG) tagged V_1_ subunit C (Vma5-mNG) in wild type (WT), *oxr1*Δ^24^, and *rtc5*Δ strains. The strains were grown to exponential phase and then deprived of glucose for 15 min before imaging. For visualization, vacuoles were stained with the lipophilic dye FM4-64. As previously reported, whereas glucose starvation in the wild type strain causes rapid V-ATPase disassembly that is accompanied by release of Vma5-mNG into the cytosol, Vma5-mNG remains on the vacuole in the *oxr1*Δ strain^24^ (**Fig. 3, upper and middle panels, respectively**). Similar to wild type, but in contrast to *oxr1*Δ, the microscopy analysis of the *rtc5*Δ cells shows diffused Vma5-mNG fluorescence throughout the cytoplasm, suggesting that Rtc5p is not required for glucose signal mediated disassembly of V-ATPase under the experimental conditions (**Fig. 3, lower panels**), an observation that is not surprising given that Oxr1p is still active in the *rtc5*Δ strain, but also consistent with the *in vitro* experiments with purified vacuoles and recombinant Rtc5p described above (**Fig. 2**).

**Figure 3.**
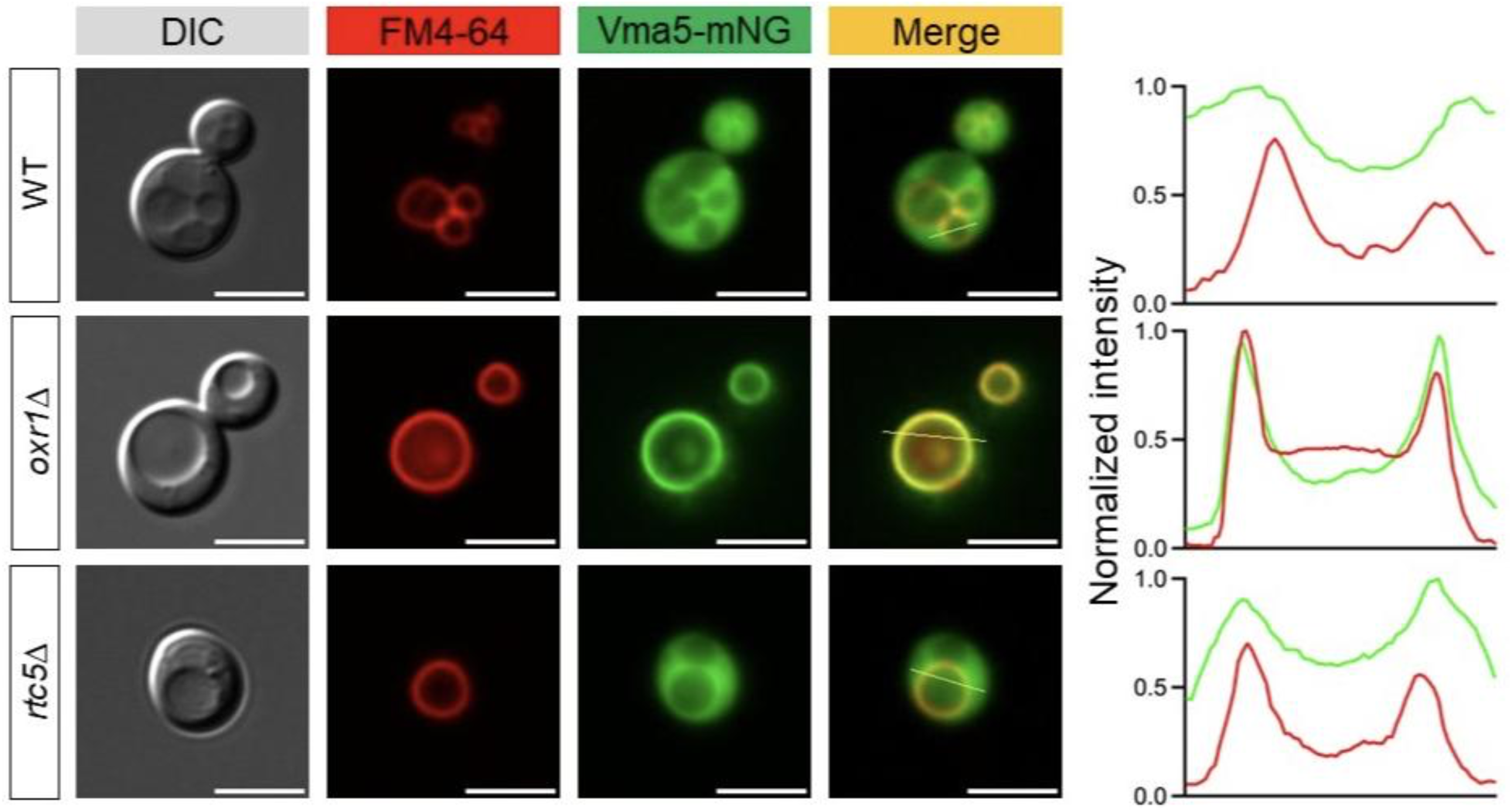
Rtc5p is not required for glucose starvation-induced V-ATPase disassembly *in vivo*. Yeast cells expressing C-terminally mNeonGreen (mNG) tagged subunit C (Vma5-mNG) in wild type, *oxr1*Δ and *rtc5*Δ strains were grown in glucose-containing media to exponential phase before depriving of glucose for 15 min. Cells were then imaged by fluorescence microscopy. Vacuolar membranes were stained with the lipophilic dye, FM4-64. 50-100 cells from two biological replicates were analyzed. Right panel shows the line scan profile of the merged image in each condition. Scale bar: 5 µm.

### Rtc5p promotes assembly of wild type V-ATPase *in vitro*

It is well documented that purified, autoinhibited wild type V_1_ and V_o_ subcomplexes and recombinant C subunit do not spontaneously assemble into holo V-ATPase, and it has been proposed that this lack of *in vitro* assembly is due a mismatch of rotary states (V_1_ is halted in state 2 and V_o_ in state 3) and the stability of the V_1_ autoinhibited conformation^21,25,41^. Consistent with this proposed model, we previously showed that an ATPase active mutant of V_1_ can assemble with V_o_ to form an active and fully coupled V-ATPase^42^. Whether V-ATPase reassembly in the cell requires reversal of V_1_ autoinhibition, however, is not known. Prior work by Kane and coworkers showed that both *in vivo* biosynthetic V-ATPase assembly and reassembly after starvation-induced disassembly requires the assembly chaperone *RAVE*^43^. According to the current model, RAVE returns cytosolic V_1_ and the C subunit to the vacuolar membrane for reassembly with V_o_ in a glucose dependent manner^28^. A recent cryoEM study showed RAVE bound to a subcomplex of V_1_ halted in rotary state 3^44^, which matches the rotary state of autoinhibited V_o_. Biochemical experiments with purified components, however, showed that while RAVE accelerated assembly of our ATPase active mutant V_1_ with V_o_, mixing RAVE bound wild type V_1_ with V_o_ did not result in functional V-ATPase^29^, suggesting that additional factors are required. Given our finding that Rtc5p does not promote holoenzyme disassembly *in vitro*, we wondered whether Rtc5p might instead play a role in the assembly process. To test this possibility, we purified autoinhibited wild type V_1_, V_o_ (reconstituted into lipid nanodisc; V_o_ND), and recombinant subunit C (**Fig. 4A**). Purified components were then mixed and incubated in absence and presence of recombinant Rtc5p at room temperature for 2 h or overnight before measuring ATPase activity (**Fig. 4B**). As the wild type V_1_ subcomplex is autoinhibited (does not hydrolyze MgATP), ATPase activity of the reaction mixtures would only be observed if the purified components (V_1_, V_o_ND and C subunit) assemble to form an active V-ATPase holoenzyme. Strikingly, whereas the sample incubated in absence of Rtc5p had virtually no ATPase activity, samples incubated in presence of Rtc5p showed a robust rate of ATP hydrolysis (9 ± 3 μmol/(mg×min) after 16 h incubation), which was >95% sensitive to the V-ATPase specific inhibitor, Concanamycin A (ConA), indicating a fully coupled holoenzyme (**Fig. 4C,D**). To confirm the formation of holoenzyme in presence of Rtc5p, we examined the post 16 h incubation mixture by negative stain transmission electron microscopy (TEM). The analysis shows the presence of holoenzyme complexes characterized by their dumbbell shape (**Fig. 4E, right panel, white circles**). In contrast, the reconstitution mixture without Rtc5p produced few (if any) holoenzyme complexes as expected from the lack of ATPase activity (**Fig. 4E, left panel**). To test whether Rtc5p can also mediate assembly of purified wild type V_1_ with V_o_ on vacuoles, we purified vacuoles from a yeast strain that has intact V_o_ on the vacuolar membrane but lacks the V_1_ subcomplex (*vma1*Δ). Vacuoles were then mixed with purified, autoinhibited, wild type V_1_ and recombinant subunit C with or without Rtc5p. After 2 h of incubation, only the sample that included Rtc5p showed significant ConA sensitive ATPase activity (0.66 ± 0.13 μmol/(mg×min)), indicating Rtc5p mediated assembly of V_1_ with endogenous V_o_ on vacuolar membranes (**Fig. 4F**). To test whether Rtc5p acts catalytically, or whether it remains bound to assembled V-ATPase, we set up a large scale (∼1 mg protein) *in vitro* reconstitution followed by SEC (**Fig. 4G**). Coomassie Blue and silver stained SDS-PAGE of the SEC elution fractions showed the presence of all V-ATPase subunits plus a band for Rtc5p running just below the A subunit, indicating that Rtc5p remained bound to the holoenzyme after facilitating its assembly.

**Figure 4.**
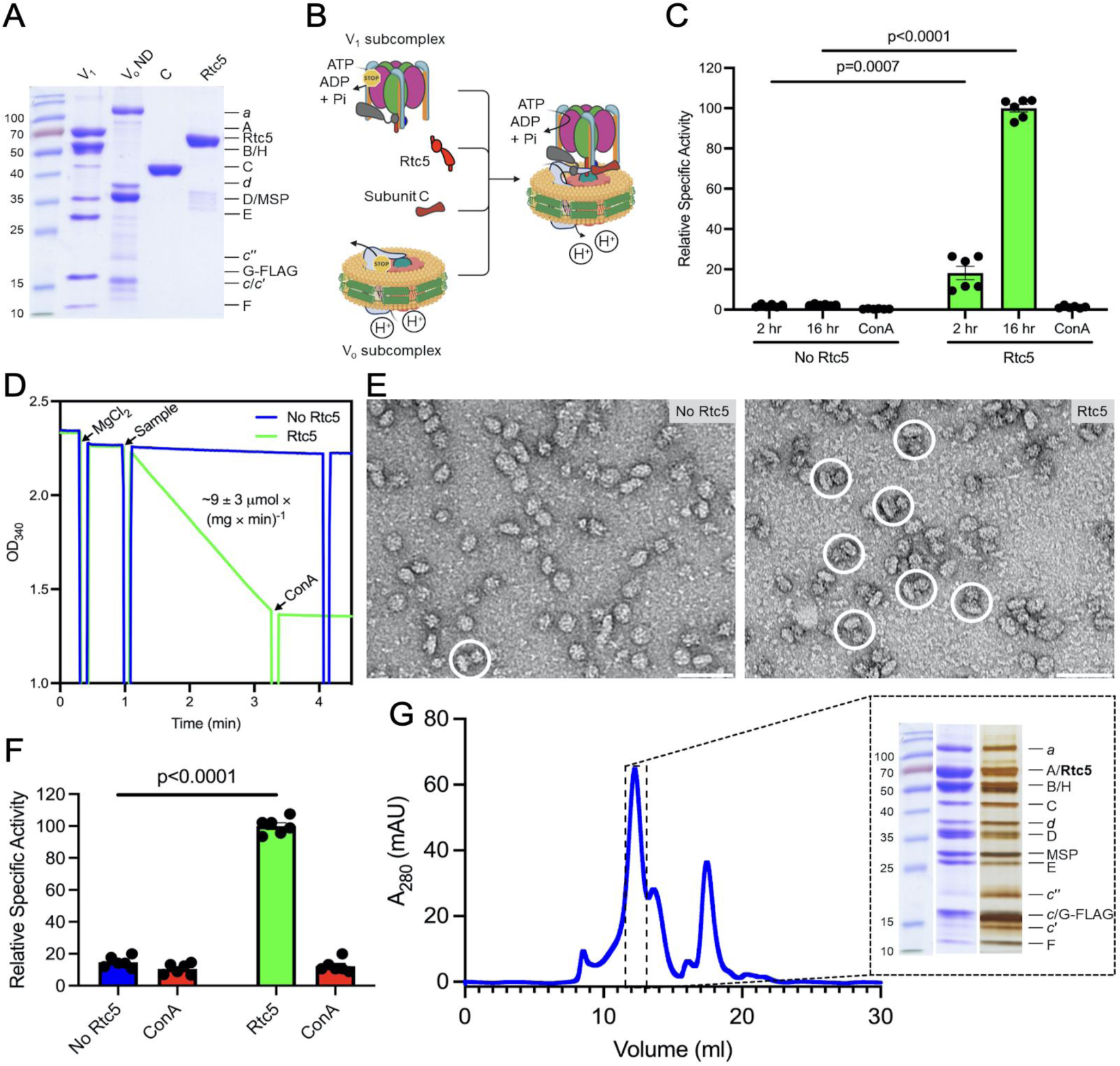
Rtc5p assembles wild type V-ATPase in vitro. **A.** Coomassie Blue stained SDS-PAGE gel of purified wild type V_1_, lipid nanodisc reconstituted V_o_ (V_o_ND), recombinant C and Rtc5p. **B.** Schematic of Rtc5p mediated *in vitro* assembly of V-ATPase. **C.** Purified wild type V_1_, V_o_ND and recombinant C were incubated at a molar ratio of 1:1:3 with or without Rtc5p in a 3:1 ratio over V_1_ for up to 16 h at room temperature. ATPase activity was measured after 2 and 16 h of incubation. ConA sensitivity of the reaction mixtures post 16 h incubation is shown. Individual data points of six tests from two biological preparations are shown. Data are presented as mean ± SEM. **D.** Representative ATPase assay traces of the post 16 h reaction mixtures from panel (C). ATPase activity is measured in an ATP regenerating assay in which the rate of ATP hydrolysis is proportional to decrease of the absorbance of NADH at 340 nm. The average of six measurements from two biological preparations gave a specific activity of ∼9 ± 3 μmol/(mg×min) for the assembly mix containing Rtc5p. **E.** Negative stain EM analysis of assembly mixtures with (right) and without (left) Rtc5p after 16 hours incubation from panel (C). A representative micrograph of a total of ∼75 images collected in three separate sessions from two biological preparations is shown. Assembled V-ATPase complexes are highlighted by white circles. Scale bars are 50 nm. **F.** Purified vacuoles from *vma1*Δ strain, wild type V_1_ and recombinant C were incubated with (green) or without (blue) Rtc5p for 2 hours at room temperature before measuring ATPase activity. ConA sensitivity of the reaction mixtures is shown as a control (red bars). Individual data points of six tests from two biological preparations are shown. Data are presented as mean ± SEM. **G.** Size exclusion chromatography elution profile of *in vitro* assembled V-ATPase on a Superose 6 increase column (10 mm × 300 mm) and SDS-PAGE analysis of assembled complex.

### CryoEM structures of V-ATPase assembled with Rtc5p and Rtc5p bound V_1_ subcomplex

SDS-PAGE analysis of *in vitro* assembled V-ATPase indicated that Rtc5p remains bound to the complex following assembly. V-ATPase assembled in presence of Rtc5p is hereafter referred to as the *post-assembly* complex, V_1_V_o_ND:Rtc5^recon^ (**Fig. 4G**). In an effort to gain insight into the mechanism by which Rtc5p facilitates *in vitro* V-ATPase assembly, we determined cryoEM structures of V_1_V_o_ND:Rtc5^recon^, as well as V_1_ subcomplex and purified wild type V-ATPase vitrified in presence of added Rtc5p.

### Structure of V-ATPase assembled in presence of Rtc5p (*post-assembly* complex)

SEC purified V_1_V_o_ND:Rtc5^recon^ was concentrated and vitrified for cryoEM structure determination. 3-D classification of ∼200,000 particle images extracted from a dataset of 7819 micrographs revealed three major populations of V-ATPase complexes (**Appendix Fig. S3**). The majority of complexes (85.3%) were halted in rotary state 1, with a prominent extra density bound at the mEAK7 site (the C-terminal domain of the B subunit next to peripheral stator EG3, see **Fig. 1C**), including 22.4% that had only weak density for subunit H (**Appendix Fig. S3**). Using MultiBody and focused refinement of the best resolved class of particles that showed strong density both for subunit H and the unassigned polypeptide in the mEAK7 site (∼46%), we obtained maps of V_1_, V_o_ND, *a*_NT_:H, the extra density, and subunit C at 3.5, 4, 4, 3.9 and 4.4 Å resolution, respectively (**Fig. 5A**; **Appendix Fig. S3**). As there is no structure available for Rtc5p, we used the model available from the AlphaFold^45^ server (**Fig. 5B**). Sequence analysis indicates that Rtc5p has an N-terminal myristoylation motif followed by an EF-hand like fold, a canonical TLDc domain, and a C-terminal segment that’s predicted to be mostly a helical (CTH) (**Fig. 1B**). Whereas AlphaFold’s prediction of the EF-hand and TLDc domain is with high confidence, there is uncertainty as to the position of the C-terminal a helix (**Fig. 5B**). Fitting the AlphaFold model of Rtc5p into the EM density revealed a close to perfect match for the EF-hand and TLDc domain (**Fig. 5C**). Strikingly, the C-terminal a helical segment is seen to insert deep into the catalytic hexamer, making contact with the central rotor (subunits D,F) and the neighboring B subunit before reemerging through the catalytic interface between peripheral stators EG1 and EG2 (**Fig. 5D; Appendix Movie S1**). The 3.9 Å resolution map of Rtc5p obtained by MultiBody refinement allowed the positioning of bulky amino acid side chains for most of the model (**Fig. 5E**). Prior structural studies of bacterial V-like and eukaryotic V-type ATPases showed that two of the three catalytic AB interfaces are usually closed and the third one open, with ADP bound in one of the closed sites^46,47^. In the Rtc5p bound state 1 complex, Rtc5p is attached to the B subunit that constitutes the closed AB interface containing ADP (**Fig. 5F**). As a consequence of the insertion of Rtc5p’s C-terminal a helix, however, the C-terminal domain of the B subunit that constitutes the closed but empty catalytic site is pushed aside, thereby partially opening that site (**Fig 5G**). At the same time, the central rotor is bent and slightly twisted to accommodate the partial opening of the normally closed site (**Appendix Movie S2**). In summary, Rtc5p binding to the V-ATPase involves insertion of Rtc5p’s C-terminal a helix into the catalytic hexamer, with concomitant partial opening of a normally closed catalytic AB interface.

**Figure 5.**
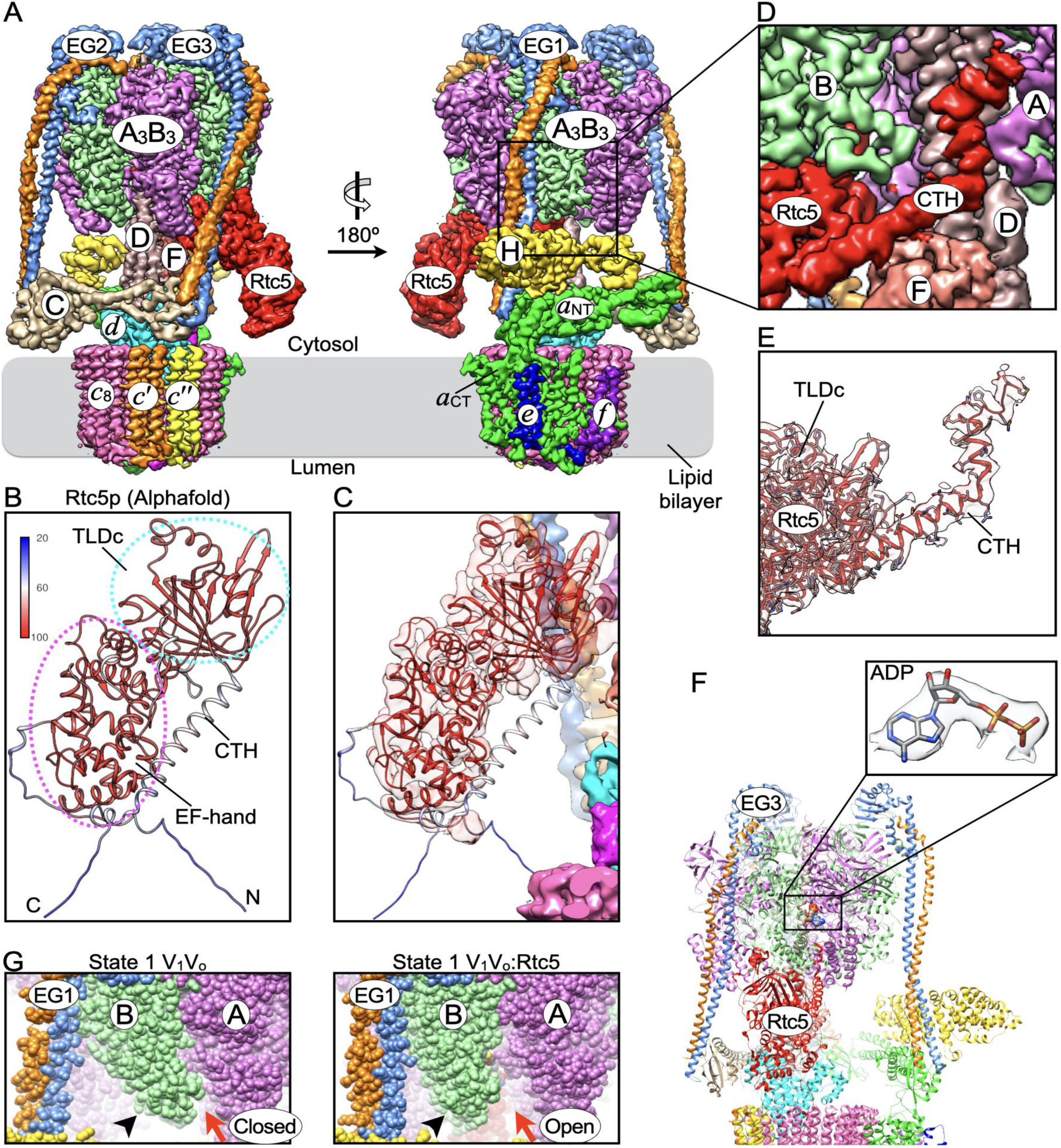
CryoEM structure of yeast V-ATPase assembled in presence of Rtc5p. **A.** CryoEM reconstruction (composite map) of V-ATPase assembled in presence of Rtc5p. **B.** AlphaFold model of yeast Rtc5p. Model confidence as provided by AlphaFold is indicated by the color key, with high to low confidence going from red to blue. **C.** Fit of the Rtc5p AlphaFold model into the cryoEM density showing a good match for the EF-hand and TLDc domain. **D.** Rtc5p’s C-terminal a helical segment inserts into the A_3_B_3_ catalytic core. For clarity, subunit H and the front-facing AB pair are not shown. **E.** The EM density for Rtc5p was refined to a resolution of 3.9 Å, allowing fitting of bulky side chains for most of the polypeptide. **F.** View of the Rtc5p bound V-ATPase rotated 90° from the view shown on the right in panel (**A**). Rtc5p is bound to the C-terminal domain of the B subunit that forms a closed catalytic site in the rotary state 1 V-ATPase complex with bound MgADP. **G.** Insertion of Rtc5p’s C-terminal a helical segment results in the partial opening of the catalytic site that’s distal to Rtc5p’s binding site.

### CryoEM structure analysis of V-ATPase vitrified in presence of added Rtc5p

The remaining ∼14.6% of the particles from above cryoEM analysis of Rtc5p-assembled V-ATPase appeared to be dominated by rotary state 2, with the extra density bound in the Oxr1 site, next to EG2 (**Appendix Fig. S3**). To further investigate the possibility that Rtc5p is able to bind the V-ATPase in both Oxr1 and mEAK7 sites, we vitrified purified wild type yeast V-ATPase after incubating the sample with an excess of recombinant Rtc5p. The cryoEM analysis of a dataset of ∼140,000 particles selected from 3561 movies revealed three major classes, with the most prominent class (42.3%) showing Rtc5p bound to the Oxr1 site in rotary state 2 V-ATPase, and the second class (33.6%) showing Rtc5p bound to the state 1 enzyme in the mEAK7 site. The third class (24.1%) did not show density for Rtc5p and was composed of a mixture of rotary states dominated by state 3 (**Appendix Fig. S4**). The Rtc5p bound state 1 and 2 complexes were refined to ∼4 Å resolution, and in both complexes, as already described for V_1_V_o_ND:Rtc5^recon^ (**Fig. 5**), Rtc5p’s CTH is seen inserted into the catalytic hexamer, partially opening the closed catalytic sites distal to Rtc5p’s binding sites (**Appendix Fig. S4, bottom panels**). A comparison of the map representing the particles in rotary state 1 with Rtc5p in the mEAK7 site (**Appendix Fig. S4**) with the map of the post-assembly complex (V_1_V_o_ND:Rtc5^recon^, **Fig. 5**) reveals a near-perfect match, except for differences in local resolution due to the different number of particle images averaged (**Appendix Fig. S5A**). While the B subunits that contribute to the Oxr1 and mEAK7 sites have the same conformation when bound to Rtc5p (backbone RMSD 0.55 Å for the entire polypeptide), the Rtc5p that’s bound in the Oxr1 site is pushed aside by ∼8 Å, likely due to the presence of the C subunit (**Appendix Fig. S5B**). We previously showed that binding of Oxr1p to state 1 V-ATPase leads to a repositioning of the C subunit and the N-terminal domain of peripheral stator EG2 (**Appendix Fig. S5C**), a conformational change that results in a weakening (and eventual breaking) of the interface between *a*_NT_ and the C subunit foot domain (C_foot_). Such a rearrangement of C and EG2 is not seen with Rtc5p bound in the Oxr1 site (**Appendix Fig. S5D**), which might be one reason why Rtc5p binding to V-ATPase does not lead to enzyme disassembly. In summary, the cryoEM data show that Rtc5p binds holo V-ATPase in a rotary state dependent manner, with Rtc5p occupying the Oxr1 site in rotary state 2 and the mEAK7 site in state 1.

### CryoEM structure of Rtc5p bound V_1_-ATPase (*pre-assembly* complex)

When purified from the yeast cytosol, wild type V_1_ is autoinhibited and predominantly halted in rotary state 2, with a molecule of ADP bound in the catalytic site next to the Oxr1 site^41,47^. Since Rtc5p can bind intact V-ATPase in both rotary states 1 and 2 (**Appendix Fig. S4**), we wished to obtain a structural picture of the interaction of Rtc5p with autoinhibited wild type V_1_. We incubated purified V_1_ with recombinant Rtc5p at an equimolar ratio and vitrified the sample for cryoEM structure determination. 3-D classification of a dataset of ∼142,000 particle images extracted from 2047 micrographs revealed three major classes of V_1_ complexes (**Appendix Fig. S6**). Approximately 16.7% of the particles represented a V_1_ complex previously referred to as *complete V_1_*^41^ as it contains subunit C, which is seen wedged between subunit H and the central DF rotor (**Appendix Fig. S6, bottom right panels**). In the V_1_ subcomplex, the rotary state is determined based on the position of the central rotor relative to the single copy inhibitory subunit H, and as reported before^41^, complete V_1_ is halted in rotary state 2, with the open catalytic site above the H subunit (**Appendix Fig. S6, bottom right panels**). The largest class of particles (∼46.9%) did not show density for the C subunit, but an Rtc5p shaped density bound in the Oxr1p site at the C-terminal domain of the B subunit located next to peripheral stator EG2 (**Fig. 6A**). When refined to a resolution of ∼3.7 Å, the map of Rtc5p bound V_1_ displayed virtually no density for the H subunit, suggesting that H is either missing or highly flexible. A low-pass filtered version of the 3.7 Å map, however, revealed clear density for H (**Fig. 6B**), indicating that while H is present, the subunit is highly dynamic in the V_1_:Rtc5p complex. Based on the position of subunit H, the V_1_:Rtc5p complex is also in rotary state 2. Thus, while Rtc5p is bound in the Oxr1p site, the Rtc5p bound complex is halted in rotary state 2 instead of state 1 as seen previously for Oxr1p bound V_1_(C)^15^. As in the intact V-ATPase assembled *in vitro* (**Fig. 5D**), Rtc5p’s C-terminal a helical segment is seen inserted into the catalytic hexamer in the V_1_:Rtc5p complex (**Fig. 6C**), with the insertion partially opening the catalytic site distal to Rtc5p’s binding site that’s normally closed (**Fig. 6C, see red arrowhead in right panel**). Moreover, as in the holoenzyme, the catalytic AB pair next to Rtc5p’s binding site is closed and has ADP bound (**Fig. 6C,D**). The state 2 V_1_:Rtc5p complex is hereafter referred to as the *pre-assembly complex*. A third (minor) class of particles (∼15.7%) from the V_1_ + Rtc5p dataset has two Rtc5p shaped densities bound that occupy both the Oxr1p and mEAK7 sites (**Appendix Fig. S6, bottom left panels**). The CTH of the Rtc5 molecule bound in the mEAK7 site, however, is not resolved, which suggests (i) that the catalytic hexamer can only accommodate one Rtc5p C-terminal a helix, and (ii), that the interaction of the TLDc domain with the C-terminal domain of the B subunit provides sufficient binding energy for Rtc5p to remain attached under these conditions. Of note, the complete V_1_ (**Appendix Fig. S6, bottom right panels**) does not have density for Rtc5p in either Oxr1p or mEAK7 sites, suggesting that this conformation of V_1_ is incompatible with Rtc5p binding. Overall, Rtc5p binds V_1_ in rotary state 2, and while Rtc5p can simultaneously occupy both Oxr1p and mEAK7 sites, only the Rtc5p bound to the Oxr1p site inserts its C-terminal a helix into the catalytic core.

**Figure 6.**
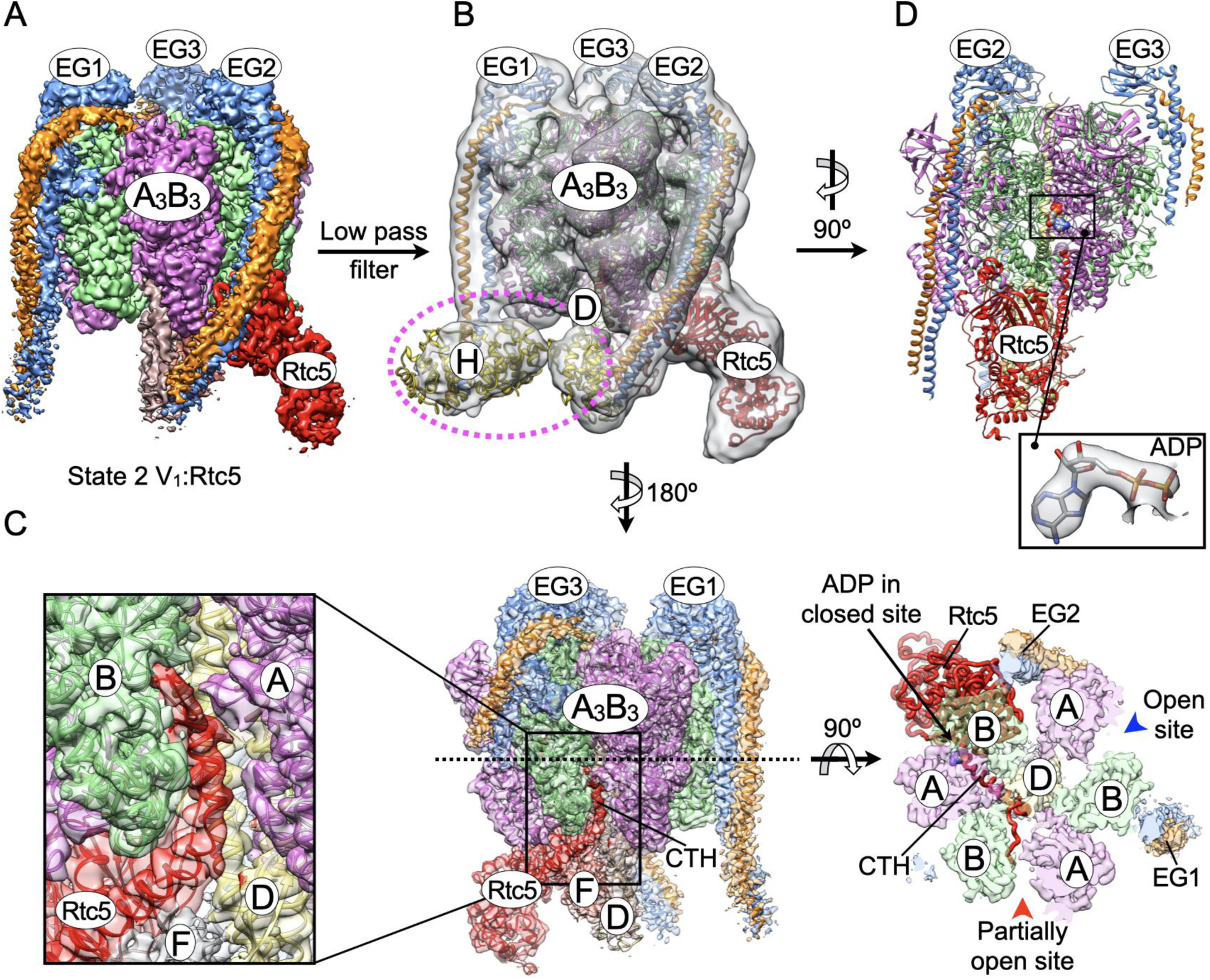
Interaction of Rtc5p with wild type V_1_ in the pre-assembly complex. **A.** 3.7 Å map of Rtc5p bound wild type V_1_. **B.** Low-pass filtered map of Rtc5p bound V_1_. The density for subunit H is highlighted by the pink dashed oval. Based on the position of subunit H, Rtc5p bound V_1_ is halted in rotary state 2. **C.** As in the Rtc5p bound holoenzyme (Fig. 5), Rtc5p inserts its CTH into the catalytic core of the V_1_ subcomplex, with Rtc5p’s very C-terminus reaching into a catalytic site distal to Rtc5p’s binding site. Left panel, zoomed in view of Rtc5p’s CTH emerging through the opened catalytic site. Right panel, view in middle panel rotated 90°, with the top-half of the molecule above the dashed line removed for clarity. The open and partially open sites are indicated by the blue and red arrowheads, respectively. ADP in the closed site next to Rtc5p is indicated by the black arrow. **D.** Rtc5p is bound to the B subunit that forms the closed catalytic site containing ADP. Inset: EM density representing bound ADP.

### Rtc5p does not activate wild type V_1_

As mentioned above, we previously showed that functional V-ATPase can be assembled from V_o_ and a mutant V_1_ subcomplex in which MgATPase activity is restored^42^. As Rtc5p binding to autoinhibited V_1_ results in a partial opening of a second catalytic site, we wondered whether the V_1_:Rtc5p pre-assembly complex regained MgATPase activity. To test this hypothesis, we incubated purified wild type V_1_ subcomplex with or without an equimolar amount of Rtc5p for 2 h at room temperature and measured ATPase activity. None of the samples, however, showed a significant rate of ATP hydrolysis, suggesting that Rtc5p does not activate wild type V_1_ under the experimental conditions (**Appendix Figure S7, top left and middle panel**). As Rtc5p alone did not activate autoinhibited V_1_, we repeated the experiment after adding either equimolar recombinant subunit C (Vma5p), or 4 mM MgATP, or both. Of note, MgATP and subunit C had been shown previously to enhance the ability of Oxr1p to inhibit holoenzyme and ATPase active mutant V_1_-subcomplex, respectively^15^. With Rtc5p, however, none of the additions tested had any effect on V_1_’s ability to hydrolyze bulk ATP (**Appendix Figure S7, top right and bottom left and middle panels**). In summary, the experiments indicated that Rtc5p does not reverse the autoinhibition of wild type V_1_, indicating that Rtc5p-mediated assembly does not require restoration of V_1_’s ATP hydrolytic activity.

### ATP disrupts the Rtc5p-V_1_ interaction, thereby blocking *in vitro* holoenzyme assembly

In cells, V-ATPase disassembly and/or reassembly occurs on a timescale of a few minutes^22,48^. The above experiments showed that Rtc5p-mediated V-ATPase assembly is a slow process, and while binding to autoinhibited V_1_ opens a second catalytic site, this conformational change did not result in bulk ATP hydrolysis activity. Previously, we have shown that the Oxr1p mediated *in vitro* disassembly process is also slow, but that the process reaches the physiological rate upon addition of ATP^24^. We therefore asked whether ATP binding to the newly opened catalytic site could make the *in vitro* assembly process faster. To answer this, we repeated the Rtc5p mediated assembly in the presence of ATP (**Fig. 7**). Surprisingly, presence of the nucleotide resulted in close to complete inhibition of Rtc5p-mediated holoenzyme assembly as evident from ATPase activity measurements (**Fig. 7A**) and negative stain TEM (**Fig. 7B, upper panel**). The cryoEM structure of Rtc5p bound V_1_ shows one open and one partially open catalytic site (**Fig. 6C**). Since wild type V_1_ purified from yeast has only one molecule of ADP bound in the closed site^41,47^, it can be assumed that the fully open site above the H subunit has low affinity for ATP. This suggests that ATP binding to the site partially opened by the interaction with Rtc5p interferes with the insertion of Rtc5p’s CTH into the catalytic hexamer. To test whether the CTH is required for mediating holoenzyme formation, we deleted it to generate Rtc5pΔCTH (**Appendix Fig. S1C,D**) and carried out assembly experiments. Measurement of ATPase activity and analysis by negative stain TEM show lower activity and fewer number of holoenzymes as compared to assembly mediated by full length Rtc5p (**Fig. 7A,B**, **Fig. 4E right panel**). This suggests that while Rtc5p’s C-terminal segment is not strictly required for mediating V-ATPase assembly, its presence makes the process more efficient.

**Figure 7.**
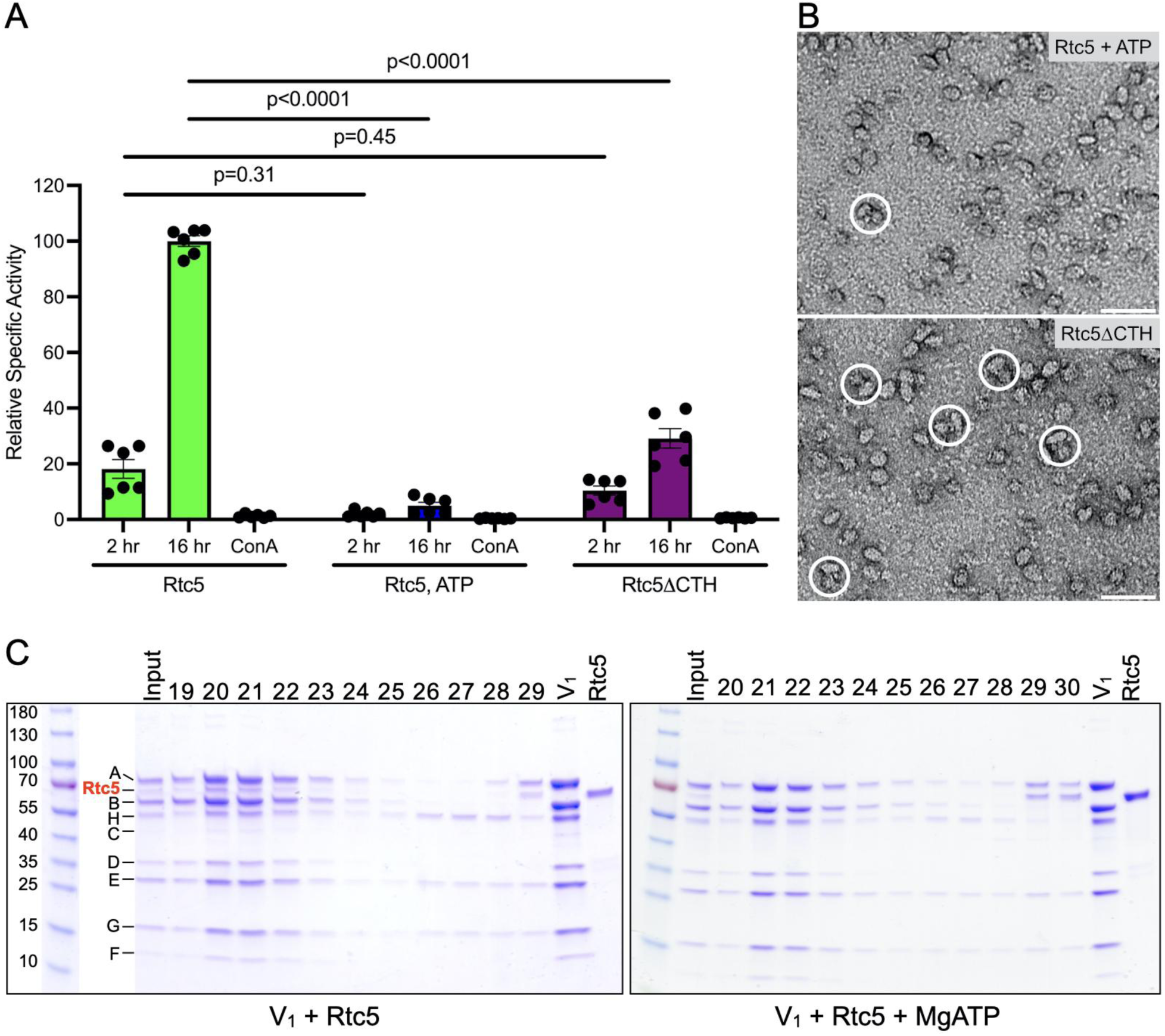
Effect of ATP and Rtc5p’s CTH on Rtc5p binding and V-ATPase assembly. **A.** Purified wild type V_1_, V_o_ND and recombinant C were incubated at a molar ratio of 1:1:3 in presence of a three-fold molar excess of Rtc5p (green), Rtc5p plus 4mM MgATP (blue), and Rtc5pΔCTH (purple) for up to 16 h at room temperature. ATPase activity was measured after 2 and 16 h of incubation. ConA sensitivity was measured at the 16 h time point. Individual data points of six tests from two biological preparations are shown. Data are presented as mean ± SEM. **B.** Negative stain EM analysis of assembly mixtures after 16 h incubation from panel (A) for the indicated conditions. Assembled V-ATPase is highlighted by the white circles. A representative of ∼75 images collected in three separate sessions from two biological preparations is shown. Scale bar: 50 nm. **C.** Coomassie stained SDS-PAGE gels of size exclusion chromatography elution fractions of mixtures of purified V_1_ + Rtc5p (left panel) or V_1_ + Rtc5p + 2 mM MgATP (right panel). Elution fraction numbers are indicated at the top of the gels.

To distinguish whether ATP binding destabilized Rtc5p binding to V_1_, or whether it interfered with the conformational changes in the V_1_:Rtc5p pre-assembly complex that are required for binding V_o_, we subjected mixtures of purified wild type V_1_ and recombinant Rtc5p with and without ATP to size exclusion chromatography (**Fig. 7C; Appendix Fig. S8**). The data show that when Rtc5p and V_1_ are mixed in absence of ATP, both components co-elute as a complex (**Fig. 7C, left panel**). When ATP is added to the sample prior to loading the column, however, the two components elute as separate species (**Fig. 7C, right panel**). This suggests that ATP binding to the catalytic site opened by Rtc5p outcompetes insertion of Rtc5p’s CTH, and that the affinity between V_1_-B and Rtc5p’s TLDc domain is too low for Rtc5p to remain bound during gel filtration.

Previously, we and others have shown that ATP hydrolysis by TLDc protein bound V-ATPase or disassembly intermediate results in TLDc protein dissociation^24,38^. While the data presented above demonstrate that Rtc5p cannot bind productively to the V_1_ subcomplex in presence of added nucleotide, we wanted to know whether ATP also destabilized binding of Rtc5p in the post-assembly complex (V_1_V_o_ND:Rtc5^recon^) (**Fig. 5**). To that end, we vitrified V_1_V_o_ND:Rtc5^recon^ with and without 600 µM MgATP for cryoEM analysis. As before (**Fig. 5**), V_1_V_o_ND:Rtc5^recon^ vitrified in absence of ATP showed Rtc5p bound in the mEAK7 site in rotary state 1 (**Appendix Fig. S9, left panel**). Analysis of the dataset collected from the sample containing ATP, however, revealed that while approximately half of the particles remained in rotary state 1 with Rtc5p bound to the mEAK7 site, the other half of the particles did not show density corresponding to Rtc5p and were about equally distributed between rotary states 1 and 2 (**Appendix Fig. S9, right panel**). Overall, the experiments show that ATP outcompetes Rtc5p binding to V_1_ and V_1_V_o_, thereby inhibiting Rtc5p mediated *in vitro* V-ATPase assembly.

### Rtc5 localizes to the vacuole in response to glucose signaling

In yeast, localization of the V_1_ subcomplex and the C subunit changes depending on the glucose signal^19^. In presence of glucose, both components are localized on the vacuole as part of the holoenzyme, and they become cytosolic in response to glucose deprivation. To analyze whether Rtc5p also responds in a similar fashion, we tagged the C-terminal domain of Rtc5p with mNeonGreen (mNG) and tested its cellular localization in different growth conditions using fluorescence microscopy. The images show that in presence of glucose, the bulk of Rtc5p-mNG is on the vacuole, which is stained here with the vacuole specific dye FM4-64 (**Appendix Fig. S10A**). However, when cells are deprived of glucose, vacuolar enrichment of Rtc5p-mNG is lost, but returns to the vacuolar membrane after re-addition of glucose to the starved cells. Previously, it was shown that RAVE carries the V_1_ subcomplex and the C subunit from the cytosol to the vacuole in response to glucose re-addition to the starved cells^28^. As the glucose signal mediated change in localization of Rtc5p is similar to V_1_ subunits, we asked if RAVE is also required to bring Rtc5p to the vacuole under these conditions. To answer this, we deleted *RAV1*, a core subunit of the RAVE complex, from the Rtc5p-mNG strain (*rav1*Δ) and collected fluorescence images under different glucose conditions. The analysis showed that Rtc5p-mNG still responds to the glucose signal in the *rav1*Δ strain, suggesting that the RAVE complex is not required for Rtc5p’s recruitment to the vacuole (**Appendix Fig. S10B**).

### Multiple pathways may exist for *in vivo* reassembly

To investigate Rtc5p’s putative *in vivo* role in V-ATPase (re)assembly, we expressed subunit C with a C-terminal mNeonGreen tag (C-mNG) in wild type^24^ and a yeast strain deleted for Rtc5p (*rtc5*Δ). Fluorescence microscopy of cells under these growth conditions shows that in both wild type and *rtc5*Δ strains, C-mNG is localized on vacuoles in glucose fed conditions, becomes cytosolic in response to glucose withdrawal, and returns to the vacuole upon re-addition of glucose to the starved cells (**Fig. 8A,B**). This suggests that even in absence of Rtc5p, C-mNG can travel back to the vacuole for reassembly, which is not unexpected given that RAVE is still functional in this strain. As stated above, RAVE carries the V_1_ subcomplex to vacuoles in response to glucose-fed or re-add condition. To investigate whether RAVE is carrying V_1_ and/or the C subunit to the vacuole during reassembly in the *rtc5*Δ strain, we deleted *RAV1* from both strains and carried out fluorescence microscopy as before. The images show that in both *rav1*Δ and *rtc5*Δ*rav1*Δ strains, Vma5-mNG is cytosolic in all conditions (**Appendix Fig. S11A,B**) indicating that RAVE is carrying subunit C (and V_1_) back to the vacuole in the *rtc5*Δ strain. While it is well established that RAVE delivers V_1_ and C to the vacuole, an *in vitro* study suggested that RAVE cannot catalyze the reassembly process itself^29^. We and Klössel *et al*.^33^ found that Rtc5p deletion did not produce a classic V-ATPase mutant, or Vma^-^, phenotype (**Appendix Fig. S2**), but as manifestation of the phenotype requires enzyme activity to be reduced to less than 30%, we therefore asked whether assembly of functional V-ATPase is impaired in the *rtc5*Δ strain. To answer this question, we purified vacuoles from wild type and *rtc5*Δ strains and found that while V-ATPase is functional in the *rtc5*Δ strain, vacuoles from the *RTC5* deletion strain have slightly less activity (∼15%) compared to wild type (**Fig. 8C**). Probing for V-ATPase on these vacuoles, however, revealed a comparable level of peripheral V_1_ subcomplex in both strains (**Fig. 8D**), suggesting that some of the V-ATPase complexes on the vacuole are improperly assembled (inactive) when Rtc5p is missing. In summary, above results indicate that while Rtc5p may play a role in ensuring proper V-ATPase assembly on the vacuole, it appears that the bulk of glucose driven rapid (re)assembly is not Rtc5p’s primary role under the conditions of the *in vivo* experiments. Thus, more work will be needed to uncover Rtc5p’s exact molecular role in V-ATPase regulation by reversible disassembly.

**Figure 8.**
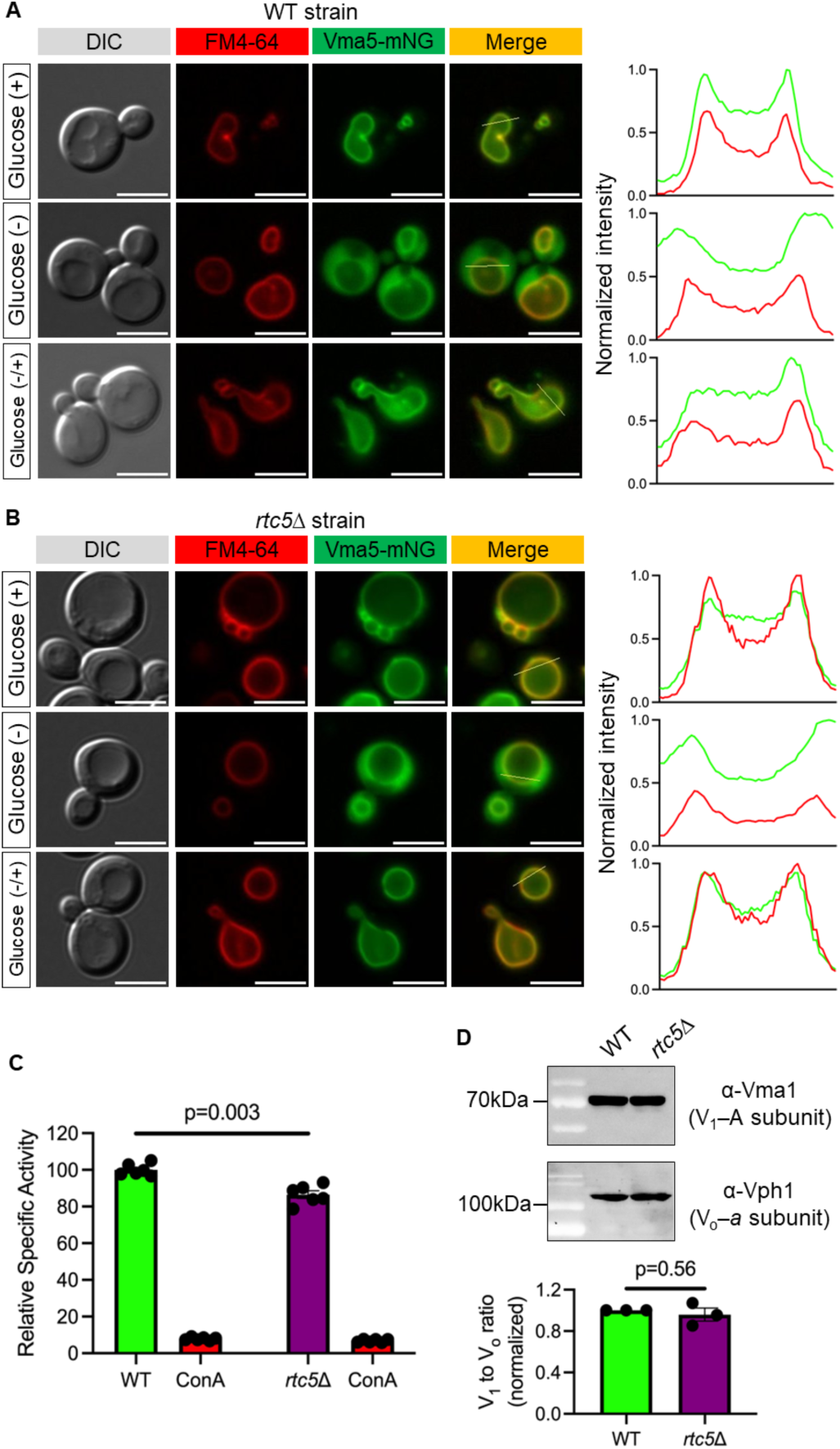
V-ATPase undergoes glucose-mediated (re)assembly in the *rtc5*Δ strain. **A-B.** Yeast cells expressing C-terminally mNG tagged subunit C (Vma5-mNG) in wild type (panel A) and *rtc5*Δ (panel B) backgrounds were grown in glucose-containing media to exponential phase before depriving of glucose for 15 min followed by 15 min of more growth after re-adding glucose. Cells were withdrawn from each growth condition, and immediately imaged by fluorescence microscopy. In both wild type and *rtc5*Δ strains, Vma5-mNG is mostly on the vacuole in glucose fed condition, gets released into the cytoplasm in response to glucose deprivation, and returns to the vacuole after re-adding glucose. Vacuoles were stained with FM4-64 dye. Around 50-100 cells from two replicates were analyzed. Right panel shows the line scan profile of the merged image in each condition. Scale bar: 5 µm. **C.** ATPase activity of purified vacuoles from wild type (green) and *rtc5*Δ (purple) strains. Vacuoles were purified in parallel and activities were measured immediately thereafter. Inhibition of V-ATPase activity by ConA was determined in each condition and plotted as an additional control. Relative activities were normalized against the wild type vacuoles that gave average specific activity of ∼6 ± 0.5 μmoles/mg/min. Individual data points of six tests from two biological preparations are shown. Data are presented as mean ± SEM. **D.** The assembly state of the V-ATPase on purified vacuoles from wild type (green) and *rtc5*Δ (purple) strains were determined by western blot using α-Vma1 (V_1_ subunit A) and α-Vph1 (V_o_ subunit *a*) antibodies. A representative of three blots from two biological preparations is shown. Band intensities were quantified, averaged, and relative ratios of V_1_ to V_o_ were normalized against wild type (WT).

## Discussion

Here we have characterized structural features and functional consequences of the interaction of Rtc5p with the yeast V-ATPase. Yeast Rtc5p (Restriction of Telomere Capping 5)^32^ belongs to the family of TLDc domain proteins, which is conserved from yeast to humans. So far, two TLDc proteins have been described in yeast, Rtc5p^33^ and Oxr1p^31^, and six in humans, OXR1, NCOA7, TLDC2, TBC1D24, mEAK7, and IFI44^49^. From studies in yeast and mammalian systems, there is mounting evidence that TLDc proteins play a role in V-ATPase regulation^15,24,33–35,50–54^. We recently showed that Oxr1p splits yeast V-ATPase into autoinhibited V_1_-ATPase and V_o_ proton channel subcomplexes in an ATP dependent manner, and that Oxr1p is essential for glucose regulated V-ATPase disassembly *in vivo*^24^. From *in vitro* experiments with purified human V-ATPase, we further showed that the human TLDc proteins, much like yeast Oxr1p, inhibit V-ATPase activity via disassembly of the complex, with the exception of mEAK7, which activates the enzyme^35^.

Rtc5p is the structural homolog of mammalian EAK7 (mEAK7), with both proteins having an N-terminal myristoylation motif followed by an EF-hand like fold, a TLDc domain, and a C-terminal a helical segment (CTH). A recent mass spectrometry and cell biological study by Klössel *et al*. concluded that yeast Rtc5p, much like Oxr1p, functions in regulated V-ATPase disassembly *in vivo*, though less efficiently than Oxr1p^33^. The molecular details of how Rtc5p carries out its putative function in the disassembly process, however, is still lacking. To further investigate the mechanism by which Rtc5p contributes to regulated V-ATPase disassembly, we expressed the protein in *E. coli* for biophysical and biochemical experiments. From ATPase activity and pelleting assays, we found that Rtc5p has no obvious impact on the enzyme’s activity, and that treatment of purified vacuoles with Rtc5p does not lead to disassembly of the complex *in vitro*. Moreover, using our standard assay where V-ATPase disassembly is induced by 15 min glucose withdrawal and visualized using mNeonGreen tagged subunit C (C-mNG), we observed that deletion of the *RTC5* gene (*rtc5*Δ) had no obvious impact on the localization of C-mNG as a function of glucose availability. While this finding was not surprising considering that Oxr1p is still present in the *rtc5*Δ strain, the overall results, however, appeared to be incongruous with Klössel *et al*.’s conclusions that Rtc5p, much like Oxr1p, functions in V-ATPase disassembly^33^. In their study, Klössel *et al*. observed more assembly and less disassembly in a strain deleted for *RTC5* as determined by comparative quantitative mass spectrometry and visualized using a C-terminal msGFP2 tag on subunit C (Vma5-msGFP2)^33^, respectively. In our experiments, on the other hand, the *in vivo* assembly state of the enzyme appeared not significantly different in the *rtc5*Δ strain compared to wild type. We currently have no explanation for this discrepancy, except to say that the two studies used different methods (quantitative mass spectrometry vs *in vitro* biochemical assays) and nutrient conditions for live-cell imaging, with the former subjecting cells to a 20 min treatment with galactose to induce V-ATPase disassembly compared to our standard 15 min glucose withdrawal. As already shown by Tabke *et al*.^55^, while galactose treatment results in dissociation of the C subunit as seen in glucose starvation, V_1_ remains on (or near) the vacuolar membrane in galactose containing media.

Since our *in vitro* experiments did not provide any evidence that Rtc5p plays a direct role in V-ATPase disassembly, we tested the opposite possibility, and to our surprise, we found that Rtc5p promoted assembly of functional holo V-ATPase from purified V_1_ and V_o_ subcomplexes *in vitro*. It is well established that enzyme disassembly, but not (re)assembly, requires ATP hydrolysis on the enzyme^22^, but that purified, autoinhibited V_1_ and V_o_ do not spontaneously assemble due to the stability of the autoinhibited conformations of the V_1_ and V_o_ subcomplexes^21^. We previously showed that assembly of functional holoenzyme can be achieved using a mutant form of the inhibitory H subunit that reverses V_1_ autoinhibition^42^. Here we find that while Rtc5p facilitates assembly *in vitro*, it does not reactivate autoinhibited V_1_ as measured in our ATP regenerating assay, and the rate of assembly is much slower compared to the rate *in vivo*^48^. Whether physiological (re)assembly involves reversing autoinhibition, however, or whether reassembly occurs via a pathway involving as of yet unseen V_1_ conformations or interactions, is not known.

From cryoEM structural studies, we found that Rtc5p can bind V-ATPase in both Oxr1p and mEAK7 sites in a rotary state dependent manner. As was seen for the interaction of mEAK7 with mammalian V-ATPase, Rtc5p is anchored to the yeast enzyme via an interaction of its TLDc domain with the C-terminal domain of a V_1_-B subunit, with the insertion of its CTH leading to partial opening of a second catalytic site. Unlike mEAK7, however, which opens the catalytic site whose B subunit it binds to, Rtc5p opens a catalytic site distal to its binding site. The reason for this difference is that Rtc5p’s significantly longer CTH can wrap around the central DF rotor to reach into the neighboring catalytic site. While Rtc5p does not reactivate autoinhibited V_1_ in bulk ATPase measurements, addition of ATP disrupts the Rtc5p-V_1_ interaction, suggesting that ATP binding, possibly at a newly opened catalytic site, alters the affinity of the interaction. Consistent with this observation, and equally puzzling considering the abundance of ATP in the living cell, addition of the nucleotide to mEAK7 bound V-ATPase also leads to release of the TLDc protein from the enzyme^38^. While we did not observe activation of yeast V-ATPase with Rtc5p, we have previously shown that mEAK7 stimulates human V-ATPase activity when added to the enzyme engaged in ATP hydrolysis^35^, indicating that these proteins may bind transiently during turnover to regulate enzyme activity. mEAK7, which is predominantly seen anchored to the lysosome via its N-terminal myristoyl group, has been implicated in activating of an alternative mTOR signaling pathway^56^. Whether Rtc5p is involved in TOR signaling in yeast, however, is not known.

What is the mechanism of Rtc5p mediated holoenzyme assembly? The cryoEM structure of the pre-assembly complex shows the majority of particles with Rtc5p bound to the Oxr1 site in rotary state 2. In the post-assembly complex, however, Rtc5p is predominantly bound to the mEAK7 site in the state 1 holoenzyme. In both cases, a molecule of ADP is seen in the closed catalytic site whose B subunit is in contact with Rtc5p’s TLDc domain. One possibility is that during assembly, Rtc5p and the ADP molecule dissociate from the Oxr1 site and rebind at the mEAK7 site. In this scenario, the complex would change rotary states from 2 (pre-assembly complex) to 3 (possibly a transient assembly intermediate), where V_1_ would be able to bind state 3 V_o_. As we have shown, Rtc5p does not bind the state 3 enzyme, likely due to steric hindrance with subunit H. Therefore, once the pre-assembly complex visits state 3 and binds V_o_, Rtc5p gets released. The complex would then have to change to the more stable rotary state 1 so that Rtc5p can rebind at the mEAK7 site. In this model, Rtc5p binding would have to lower the activation barrier for the pre-assembly complex to visit state 3. Another possibility is that subunit H gets transiently released from the pre-assembly complex to ease V_1_’s transition to state 3. Once the complex assembles, H would rebind at a different peripheral stator. In this mechanism, Rtc5p and ADP could stay bound during assembly as no net rotation of the central rotor relative to Rtc5p would be required. Support for the latter model comes from the observation that H binding in the pre-assembly complex appears to be highly dynamic. Determining whether Rtc5p mediated assembly occurs according to one of the two models described above, or a completely different mechanism, however, will require further study.

## Materials and Methods

### Plasmid construction

Plasmids pET28a_His_Rtc5 and pE28a_His_Rtc5ΔCTH were generated using restriction free cloning as described previously^24,57^. Briefly, the gene encoding full length Rtc5p was amplified from yeast genomic DNA using Rtc5_pET_F (5′-GAG ATA TAC CAT GGG TCA TCA TCA TCA TCA TCA TCA CGG ACA GTC TTC TTC AAT CAG TTC-3′) and Rtc5_pET_R (5′-GTG CTC GAG TGC GGC CGC AAG CTC ACA TTG AAC CGC CAC C-3′) primers pair, and the PCR product was used as megaprimer to insert into a modified pET28a vector^58^ to construct the pET28a_His_Rtc5 plasmid. To generate the pET28a_His_Rtc5ΔCTH plasmid, the gene for Rtc5p without the C-terminal helix (Δ516-567 a.a.) was PCR amplified using pRtc5_pET_F and Rtc5_CTD_pET_R (5′-GTG CTC GAG TGC GGC CGC AAG CTT AAC CAC AGC CCC ATA CCT C-3′) primers before inserting into the pET28a vector. In both cases, insertion was verified by restriction digestion and DNA sequencing.

### Generation of yeast strains

All yeast strains generated in this study (**Appendix Table S2**) were generated in the 5Aα parental background using a standard lithium acetate protocol. Briefly, a specific gene was PCR amplified and transformed into the target yeast strain for genomic integration by homologous recombination. Yeast strains containing the correct insert were then selected on drug containing YEPD plates and confirmed by colony PCR (Phire Plant Direct PCR Master Mix; ThermoFisher, catalog no. F160S) and DNA sequencing.

### Expression and purification of Rtc5p and Rtc5pΔCTH

N-terminally 7×His tagged Rtc5p and Rtc5pΔCTH were expressed in BL21DE3 *E. coli* strains transformed with pET28-His-Rtc5 and pET28a-His-Rtc5ΔCTH plasmid, respectively. Cells were grown to an OD_600_ of 0.6-0.8 in RB (LB + 0.2 % glucose) + 50 µg/ml kanamycin media before induction with 0.5 mM isopropyl-β-D-1-thiogalactopyranoside (IPTG) for 16-18 hours at 20 °C. Cells were harvested by centrifugation at 4,000 × g for 30 min and stored at −20 °C until use. Cells were thawed, supplemented with 1 mM PMSF, 2 mM DTT, lysozyme (25 mg/l cells) and DNase (2 mg/l cells), and lysed by sonication. Cell debris and unbroken cells were separated by centrifugation at 13,000 × *g* for 30 min at 4 °C. The cleared lysate was loaded onto a 1 ml Ni-NTA column attached to an ÄKTA FPLC and pre-equilibrated with buffer A (20 mM Tris·HCl, pH 8.0, 250 mM NaCl, 10 mM Imidazole, 2 mM DTT). Bound protein was eluted by an imidazole gradient elution using buffer B (20 mM Tris·HCl, pH 8.0, 250 mM NaCl, 500 mM Imidazole, 2 mM DTT). Fractions were analyzed by SDS-PAGE and concentrated using a 10 kDa MWCO centrifugal concentrator before passing over a Superdex 200 column (16 mm × 500 mm) pre-equilibrated with 20 mM Tris·HCl, pH 7.0, 100 mM NaCl, 1.5 mM DTT for further polishing.

### Purification of wild type V_1_ subcomplex

Purification of N-terminally FLAG tagged G subunit containing wild type V_1_ was done using established protocols^15,42^. Briefly, cells were grown in YEPD media to an OD_600_ of ∼3.5 to allow the V_1_ to detach from the vacuole, harvested by centrifugation and stored at −80 °C until use. Cells were resuspended in TBSE buffer (20 mM Tris·HCl, pH 7.2, 150 mM NaCl, 0.5 mM EDTA) supplemented with 5 mM β-mercaptoethanol and protease inhibitors (leupeptin, pepstatin, and PMSF) and lysed by passage through a microfluidizer at 18000 psi. The cell lysate was cleared by two subsequent centrifugations at 4000 and 13,000 × *g*, respectively, before being passed over an anti-FLAG affinity column (5 ml) pre-equilibrated with TBSE. The anti-FLAG column was washed with 10 column volumes (CVs) TBSE and eluted with 25 ml of the same buffer containing 0.1 mg/ml FLAG peptide. After collecting the first fraction (two thirds of a column volume), the column was stopped for 45 min followed by collecting subsequent fractions of each 5 ml. Fractions were analyzed by SDS-PAGE and V_1_ subcomplex containing fractions pooled, concentrated to 2 ml, and passed over a Superose 6 Increase HiScale (16 mm × 400 mm) size-exclusion chromatography (SEC) column attached to an ÄKTA FPLC using 20 mM Tris·HCl, pH 7.2, 0.5 mM EDTA, 1 mM TCEP buffer. Fractions were then analyzed by SDS-PAGE, pooled and concentrated using a 50 kDa MWCO centrifugal concentrator. Concentrated protein was supplemented with 2.5 mM DTT and stored at 4 °C on ice.

### Purification of lipid nanodisc reconstituted V_o_ subcomplex

Purification and reconstitution of V_o_ into lipid nanodisc was carried out as described^59,60^. A yeast strain containing C-terminally calmodulin-binding peptide (CBP) tagged V_o_ *a* subunit (Vph1 isoform) was grown to OD_600_ of 12-15, harvested by centrifugation, washed in water, resuspended in lysis buffer (25 mM Tris·HCl, pH 7.4, 500 mM sorbitol, 2 mM CDTA), and stored at −80 °C until use. ∼320 g of cells were thawed and added into a 900 ml glass jar containing zirconium beads up to the ∼500 ml mark, and the remaining volume was filled with lysis buffer and supplemented with protease inhibitors (leupeptin, pepstatin, aprotinin, chymostatin, and PMSF). The mixture was cooled using an ice-salt mixture while mixing (“beating”) at 3,000 rpm in an inverted bead-beater. Cells were ruptured by eight cycles of beating with each cycle consisting of one-minute bursts at 20,000 rpm and seven-minutes of cooling at 3,000 rpm. The cell lysate was supplemented with 1 mM PMSF and cleared by centrifugation at 4000 and 13,000 × *g* to remove cell debris and mitochondria, respectively. Microsomal membranes were pelleted by ultracentrifugation at 200,000 × *g* for 2 h and resuspended in washing buffer (25 mM Tris·HCl, pH 7.4, 500 mM sorbitol) supplemented with protease inhibitors. Resuspended membranes were washed using a Dounce homogenizer and collected by centrifugation at 200,000 × *g* for 1 h. The protein concentration of washed membranes was determined using a modified BCA assay^59^ and membranes were stored at −80 °C until use. Membranes corresponding to ∼2 g of membrane protein were resuspended in cold lysis buffer to achieve a final concentration of 10 mg/ml. Resuspended membranes were supplemented with protease inhibitors and extracted with 0.6 mg/mg membrane protein (w/w) undecyl maltoside (UnDM) for 30 min while gently rotating at 4 °C. The mixture was then supplemented with 4 mM CaCl_2,_ and extraction was continued for an additional 30 min. Solubilized membrane protein was collected by ultracentrifugation at 100,000 × *g* for 1 h before passing over a calmodulin sepharose (CaM) column pre-equilibrated with calmodulin washing buffer (10 mM Tris·HCl, pH 8, 10 mM β-mercaptoethanol, 2 mM CaCl_2_, 0.06% UnDM) plus 150 mM NaCl. The column was washed with 100 ml of each calmodulin washing buffer with and without NaCl. Bound V_o_ subcomplex was then eluted from the column in 100 ml of 10 mM Tris·HCl, pH 8, 10 mM β-mercaptoethanol, 10 mM CDTA, 0.06% UnDM buffer. After collecting the first fraction (two thirds of a column volume), the column was stopped for 30 min followed by collecting 12 ml fractions. Fraction(s) containing the protein was concentrated to ∼2 ml using a 100 kDa MWCO centrifugal concentrator, and protein concentration was measured using the modified BCA assay. To reconstitute V_o_ into lipid nanodiscs (V_o_ND), detergent-solubilized V_o_, membrane scaffold protein (MSP1E3D1), and *E. coli* polar lipids were combined in disc forming buffer (20 mM Tris·HCl, pH 7.4, 0.5 mM CDTA, 100 mM NaCl, 1 mM DTT) at a molar ratio of 0.02:1:25 in presence of protease inhibitors and 1.5% UnDM. Reconstitution was carried out at room temperature for 1 h with gentle mixing followed by detergent removal using 0.4 g/ml Bio-Beads SM2 (Bio-Rad) for 2 h. Following separation from Bio-Beads, the reconstitution mixture was supplemented with 5 mM CaCl_2_ and passed over a 3 ml CaM column to remove empty nanodiscs. The column was eluted as before, and the V_o_ND-containing elution fraction was applied to a Superdex 200 Increase HiScale (16 mm × 400 mm) SEC column equilibrated in disc forming buffer containing 1 mM TCEP. V_o_ND-containing fractions were combined, concentrated, and stored at 4 °C on ice, or mixed with 20% glycerol, flash-frozen in liquid nitrogen, and stored at −80 °C for later use.

### Expression and purification of subunit C

The subunit C were expressed and purified following established protocols^15,61^. Briefly, *E. coli* Rosetta2 cells expressing N-terminally maltose-binding protein (MBP)-tagged C subunit were cultured at 37 °C in RB media (LB + 0.2% glucose) supplemented with 50 µg/ml carbenicillin and 34 µg/ml chloramphenicol until reaching an OD_600_ of 0.6-0.8. Protein expression was induced with 0.4 mM IPTG at 30 °C for 5.5 h. Cells were harvested by centrifugation, resuspended in amylose column buffer (ACB: 20 mM Tris·HCl, pH 7.4, 200 mM NaCl, 1 mM EDTA) and stored at −20 °C until use. Cells were treated with lysozyme (25 mg/l cells), DNase (2 mg/l cells), 1 mM PMSF and lysed by sonication. The lysate was clarified by centrifugation, and the supernatant was passed through a 20 ml amylose column pre-equilibrated with ACB. The column was washed in 10 CV ACB, and proteins were eluted with 25 ml ACB supplemented with 10 mM maltose in one fraction. Eluted protein was supplemented with 5 mM DTT and the MBP tag was cleaved with Prescission protease (200 units/ liter of cells) for 2 hours at 4 °C with gentle mixing. The protein was dialyzed overnight in 20 mM Tris·HCl, pH 6.5, 0.5 mM EDTA buffer in a cold room and then passed over a diethylaminoethyl (DEAE) column to capture and remove MBP. The flow-through of the column was dialyzed again in 20 mM Tris·HCl, pH 7, 0.5 mM EDTA buffer and concentrated before being further purified by a Superdex 75 SEC column (16 mm × 500 mm) in the same buffer. The purified proteins were analyzed by SDS-PAGE, concentrated using a 10 kDa MWCO centrifugal concentrator, and stored at −80 °C in 20% glycerol for later use.

### Expression and purification of MSP

Membrane scaffold protein (MSP) was expressed and purified from *E. coli* BL21DE3 cells as described^35^. Briefly, cells were grown at 37 °C in RB medium in presence of 50 μg/ml kanamycin. After reaching an OD_600_ of 0.5-0.6, expression of MSP1E3D1 (containing N-terminal 7×His, Avi tags, and a Prescission protease cleavage site) was induced with 0.5 mM IPTG for 4 h. Cells were harvested, and pellets were stored at −20 °C until use. Cell pellets were resuspended in lysis buffer (20 mM Tris·HCl, pH 8, 10 mM imidazole, 500 mM NaCl, and 6 M guanidine·HCl), sonicated, and cleared by centrifugation. The supernatant was applied to a 10 ml Ni-NTA column at room temperature, followed by washing with 5 CV of lysis buffer. Protein was refolded on the column with a slow overnight wash at 4 °C with 10 CV of buffer 1 (20 mM Tris·HCl, pH 8, 10 mM imidazole, 250 mM NaCl). The column was washed subsequently with the following buffers: 10 CV of buffer 2 (20 mM Tris·HCl, pH 8, 300 mM NaCl, 1% Triton X-100), 10 CV of buffer 3 (20 mM Tris·HCl, pH 8, 300 mM NaCl, 25 mM sodium cholate), and 10 CV of buffer 1. Protein was then eluted in 5 CV of 20 mM Tris·HCl, pH 8, 1 M imidazole, 250 mM NaCl. Protein-containing fractions were dialyzed overnight in 20 mM Tris·HCl, pH 8, 150 mM NaCl buffer at 4 °C, centrifuged to remove any aggregates, and stored at −80 °C.

### Expression and purification of Oxr1p

Expression and purification of N-terminally 7×His tagged Oxr1p was done as described^15^. Briefly, *E. coli* BL21DE3 cells were cultured at 37 °C in RB medium supplemented with 50 μg/ml kanamycin until reaching an OD_600_ of 0.6-0.8. Protein expression was induced by adding 0.5 mM IPTG at 20 °C for 16-18 h. The cells were harvested by centrifugation, resuspended in Buffer A (20 mM Tris·HCl, pH 8, 250 mM NaCl, 10 mM imidazole, 2 mM DTT), and stored at −20 °C. Thawed cells were treated with lysozyme and DNase in the presence of 1 mM PMSF and 2 mM DTT and lysed by sonication. Cellular debris was removed by centrifugation, and the supernatant was applied to a 1 ml Ni-NTA column pre-equilibrated with Buffer A. After washing the column with 10 CV of Buffer A, the protein was eluted using a 0-60% imidazole gradient over 30 CV Buffer B (20 mM Tris·HCl, pH 8, 250 mM NaCl, 500 mM imidazole, 2 mM DTT). Fractions containing Oxr1p were combined, concentrated to ∼2 ml, and applied to a Superdex 75 SEC column (16 mm × 500 mm) in 20 mM Tris·HCl, pH 7, 100 mM NaCl, 1.5 mM DTT. Fractions containing Oxr1p were pooled, concentrated using a 10 kDa MWCO centrifugal concentrator, and stored at −80 °C in 20% glycerol until use.

### Purification of yeast vacuoles

Yeast vacuoles were isolated as described^15,24^. Briefly, yeast cells were grown to an OD_600_ of ∼1.0 either in unbuffered or buffered (pH 5) YEPD media, harvested, and spheroplasted using zymolyase 100T treatment. Yeast spheroplasts were then broken in a Dounce homogenizer and vacuoles were collected from the top layer of a Ficoll gradient following two rounds of ultracentrifugation (71,000 × *g*, 4 °C, 40 min). Vacuoles were resuspended in 1.5 mM MES-Tris pH 7, 4.8% glycerol for immediate use, or stored at −80 °C. For biochemical analysis, thawed vacuoles were washed and resuspended in 10 mM Tris·HCl, pH 7.4, 1 mM EDTA before use.

### Assembly of wild type V_1_ subcomplex with V_o_ in vacuolar membranes

To study the effect of recombinant Rtc5p and Rtc5pΔCTH on V-ATPase assembly, purified wild type V_1_, V_o_ND and recombinant C mixed at a molar ratio of 1:1:3 and incubated for 16 hours at room temperature in presence of (1) TBSE buffer, (2) three molar fold Rtc5p, (3) Rtc5p plus 4 mM MgATP, and (4) three molar fold Rtc5pΔCTH. ATPase activities were measured after withdrawing samples after 2 and 16 h of incubation. For assembly with vacuoles, vacuoles purified from a *vma1*Δ strain were mixed with wild type V_1_ and recombinant subunit C (V_1_:vacuole = 1:1 (w/w) and V_1_:C =1:3 (molar ratio)) in absence and presence of three molar fold Rtc5p and incubated for 2 h before measuring the ATPase activity.

### ATPase assay

ATPase activity of vacuoles, (re)assembled V-ATPase and V_1_ subcomplex were measured in an ATP regenerating ATPase assay as described^15^. Briefly, 1 ml assay mixture containing 50 mM HEPES pH 7.5, 25 mM KCl, 0.5 mM NADH, 2 mM phosphoenolpyruvate, 5 mM ATP, and 30 units each of lactate dehydrogenase and pyruvate kinase was pre-warmed for 10 minutes at 37 °C. ATPase assays were then carried out in a temperature-controlled Varian Cary 100 Bio UV-VIS spectrophotometer where the change in absorbance at 340 nm over time was recorded in a Kinetics Application in the Cary WinUV software package version 3. To initiate the reaction, assays were first supplemented with 4 mM MgCl_2_ before adding 10 µg of protein into the assay. After stopping the reaction, assay traces for V_1_ samples were exported as CSV files and plotted in GraphPad Prism 9 software. For measuring the ATPase activity of vacuoles and (re)assembled V-ATPase, after adding the protein samples into the assay, enough time was given to establish a linear rate of ATP hydrolysis before adding 200 nM of the V-ATPase specific inhibitor concanamycin A (ConA). The rate of ATP hydrolysis was then determined using the same software package that was used to collect the data.

### V_1_ detachment from vacuoles analysed by western blot

The analysis was carried out as described before^24^. Briefly, purified vacuoles were incubated with (1) buffer, (2) Rtc5p at a ratio of 1:5 (w/w) to vacuoles plus 4 mM MgATP, (3) Oxr1p (at a ratio of 1:10 (w/w) to vacuoles) plus 4 mM MgATP, and (4) 4mM MgATP for 30 minutes at room temperature (a portion kept aside on ice served as *input* on the blot). Vacuole samples were then ultracentrifuged at 37,000 × *g* for 20 min at 4 °C. Pellets and supernatant (loaded as *pellet* and *sup*, respectively, on the blot) were separated and resuspended along with the *input* in a hot cracking buffer (50 mM Tris·HCl, pH 6.8, 1 mM EDTA, 8 M Urea, 5% SDS, 10 mM DTT). Samples were then separated by SDS-PAGE, transferred to low-fluorescence polyvinylidene fluoride membranes, and blocked for 1 h in TBST buffer (20 mM Tris·HCl, pH 7.5, 150 mM NaCl, 0.05% Tween-20) plus 5% milk protein with gentle shaking at room temperature. After blocking, blots were incubated with either mouse α-Vma1 (8B1F3) or mouse α-Vph1 (10D7) antibody overnight at 4 °C on a rocking platform. After washing in TBST buffer for three times each 10 min, goat **α**-mouse IgG Alexa Fluor Plus 488 (ThermoFisher, catalog no. A32723) antibody was added (1:2000) and incubated for 90 min at room temperature with gentle agitation. Blots were washed again as before, dried and imaged in a Bio-Rad ChemiDoc MP Imaging System. All images were analyzed using the Bio-Rad Image Lab software.

### Negative stain electron microscopy

The assembly state of *in vitro* assembled V-ATPase was determined by negative stain electron microscopy at SUNY Upstate Medical University’s TEM core facility as described^15,24^. Briefly, protein samples were diluted to 20-30 µg/ml in TBSE buffer before applying 5 µl samples onto glow discharged carbon-coated copper grids. After 1 min of incubation, grids were washed in Milli-Q water and stained with 5 µl of 1% uranyl acetate solution for 1 min. In each step, excess liquid from the grids was removed using Whatman filter paper. Micrographs were collected at 200,000 × magnification and 80 keV on a JEOL JEM-1400 transmission electron microscope equipped with a Gatan Orius SC1000 CCD camera.

### CryoEM grid preparation and data collection

Preparations of V_1_ and V_1_V_o_ND at concentrations of between 1-2 mg/ml in TBSE with or without equimolar amounts of recombinant Rtc5p were vitrified on C-flat or Au-flat holey grids (1.2/1.3 µm) using a Leica EM GP2 grid plunger set to 90% humidity, 6 °C, 7-9 s blot time, sensor blotting. Grids were screened locally on a JEOL JEM-2100F transmission electron microscope equipped with a Gatan 914 side entry cryo holder and OneView CMOS and K2 Summit direct detectors using SerialEM for automated data collection^62^. Larger datasets were obtained at the Hauptman Woodward Institute (Buffalo, NY) cryoEM facility on a ThermoFisher Glacios TEM equipped with a Falcon IVi camera.

### CryoEM data processing

CryoEM images were analyzed using cryoSPARC v4.4^63^ and RELION 5^64^. Movies were subjected to motion and CTF correction as implemented in RELION 5. Particles were picked using the blob picker (cryoSPARC) or Topaz^65^ (RELION) and particle datasets subjected to 2-D and 3-D classification to remove bad particle picks and identify structural and conformational heterogeneity. Consensus reconstructions for V_1_V_o_ND:Rtc5^recon^ and V_1_:Rtc5 were improved by focused and/or multi-body refinement as implemented in RELION5 using masks generated from molmaps as generated in UCSF Chimera^66^.

### Model building and refinement

Model building for V_1_V_o_ND assembled in presence of Rtc5p (V_1_V_o_ND:Rtc5^recon^) was started using our previous models of state 1 and state 2 yeast V-ATPase (PDBIDs: 7fda, 7fdb^15^) and the AlphaFold^45^ model of Rtc5p. Atomic models were placed into the cryoEM density using rigid body fitting in UCSF Chimera^66^ followed by manual building and automated real space refinement using Coot^67^, ISOLDE^68^, and Phenix^69^. For details of 3-D reconstruction, model building, and refinement, see **Appendix Table S1**.

### Fluorescence microscopy

The localization of mNeonGreen (mNG) tagged proteins in different strains were carried out using established protocols^24,28^. Briefly, cells were grown to exponential phase in synthetic complete (SC) media, stained with lipophilic dye FM4-64 (ThermoFisher, catalog no. T3166) in YEPD, and recovered and washed in SC media. DIC and fluorescence images of cells were then collected in glucose-fed (+) condition, 15 min after glucose deprivation (-) and 15 min after re-adding glucose (-/+) to the starved cells using DIC, TexasRed (Red) and GFP (Green) channels in a Zeiss fluorescence microscope. Images were imported into Fiji ImageJ software^70^ and then saved as a JPEG file after subtracting the background. Saved fluorescent images (Red and Green) were then reimported again into Fiji ImageJ and converted into 16 bit for drift correction using TurboReg plugin^71^ using Red image as a source and Green image as a target. Aligned images were then converted back into RGB color and merged to generate “merge image”. All line scans were performed using the “Color Profiler” plugin in Fiji ImageJ.

## Data availability

All data needed to evaluate the conclusions in the paper are present in the paper and/or the supplementary materials. Additional data are available from the authors upon request. The cryoEM maps and corresponding coordinate models are deposited in the Electron Microscopy Data Bank and Protein Data Bank (EMD-48491 and PDBID: 9MOY for composite map state 1 V_1_V_o_ND:Rtc5^recon^; EMD-49559 and PDBID: 9NN1 for composite map state 2 V_1_:Rtc5; EMD-70377 and PDBID: 9ODU for state 2 V_1_V_o_ND:Rtc5). Multibody and consensus maps for V_1_V_o_ND:Rtc5^recon^ are EMD-48485 (consensus), EMD-48486 (V_1_), EMD-48487 (V_o_), EMD-48488 (aNT:H), EMD-48489 (Rtc5p), EMD-48490 (subunit C), and for V_1_:Rtc5, EMD-49556 (consensus), EMD-49557 (V_1_), EMD-49558 (Rtc5p).

## Acknowledgements

CryoEM data presented in this study were collected at SUNY ESF, Syracuse, New York and at the Hauptman-Woodward Institute CryoEM Facility, Buffalo, New York. This work was supported by NIH grant GM141908 to S.W.

## Author contributions

Md.M.K.: conceptualization, formal analysis, investigation, methodology, resources, validation, visualization, writing - original draft preparation, review, and editing; R.E.: conceptualization, formal analysis, investigation, methodology, resources, validation, visualization, writing - review and editing; R.A.O.: methodology, resources, writing - review and editing; S.W.: conceptualization, formal analysis, investigation, methodology, resources, supervision, validation, writing - review and editing.

## Appendix Figures

**Appendix Figure S1.**
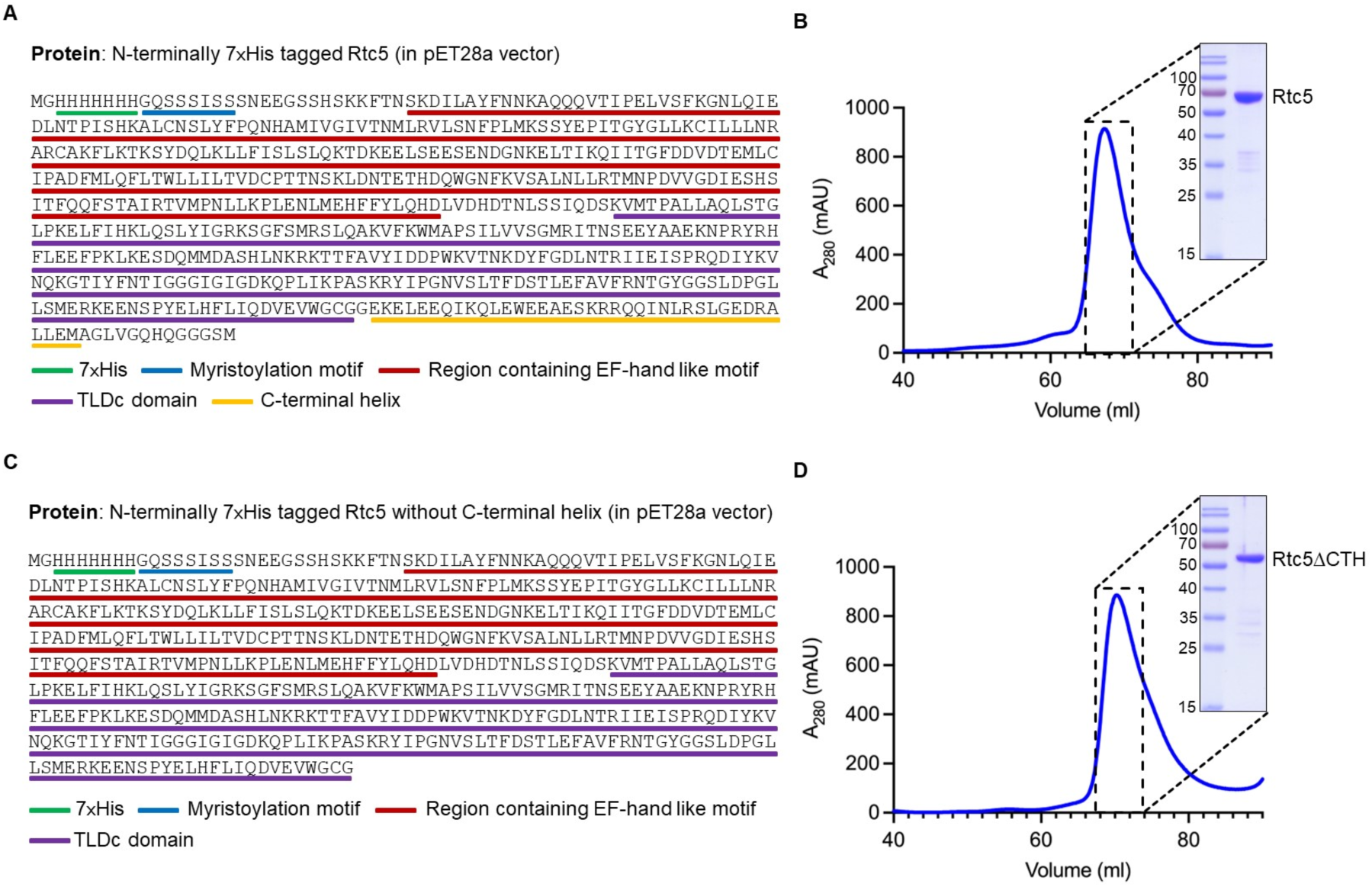
Construct design and purification of Rtc5p and Rtc5pΔCTH proteins. **A.** *RTC5* is cloned into a pET28a vector with a N-terminal 7×His tag. **B.** Size exclusion chromatography profile and SDS-PAGE of 7×His tagged full length Rtc5p. **C.** *RTC5* without its C-terminal helix (Rtc5pΔCTH) is cloned into a pET28a vector with a N-terminal 7×His tag. **D.** Size exclusion chromatography profile and SDS-PAGE of 7×His tagged Rtc5pΔCTH.

**Appendix Figure S2.**
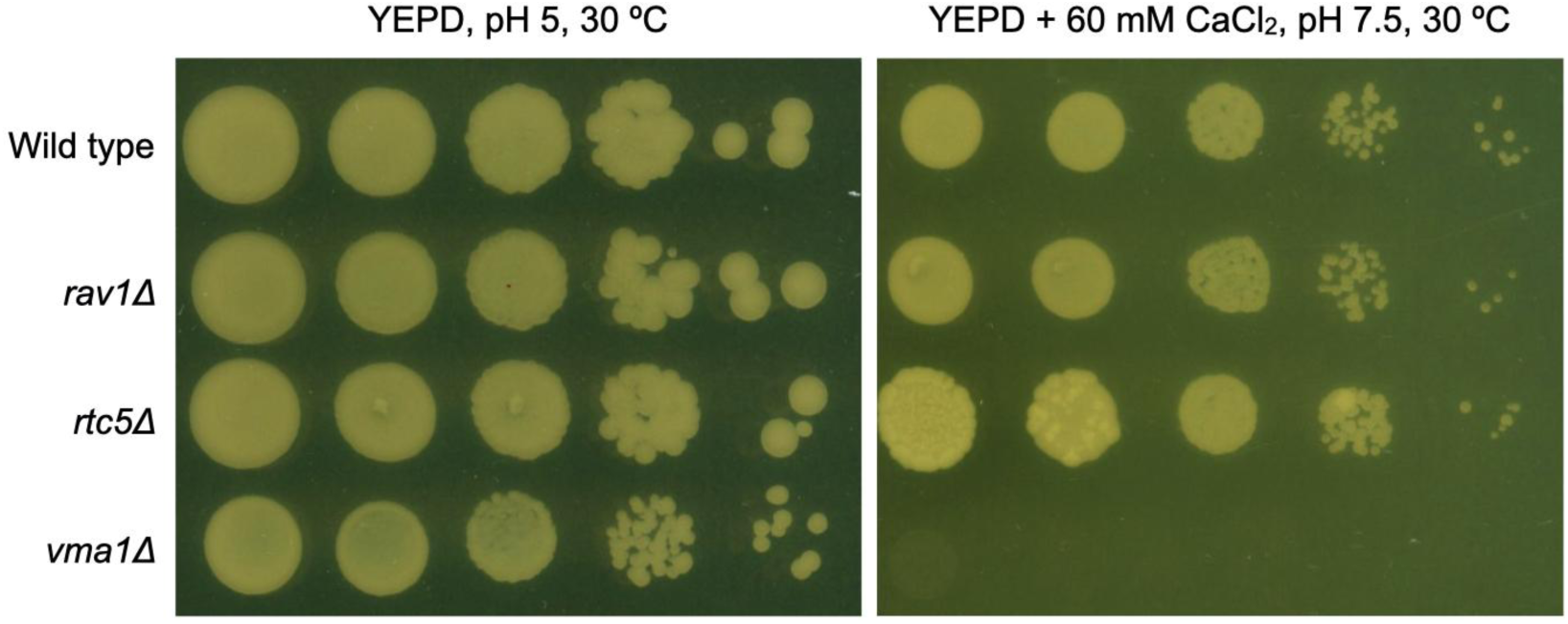
Growth phenotype of *rtc5*Δ strain. Wild type, *rav1*Δ and *rtc5*Δ cells were grown to exponential phase and serially diluted tenfold before spotting on YEPD pH 5 and YEPD pH 7.5 + Ca^2+^ plates. Plates were incubated for 3 days at 30 °C before recording images. The *vma1*Δ strain, which produces the classic Vma^−^ growth phenotype in YEPD pH 7.5 + Ca^2+^ medium, was included as control. A representative assay from a total of three repeats is shown.

**Appendix Figure S3.**
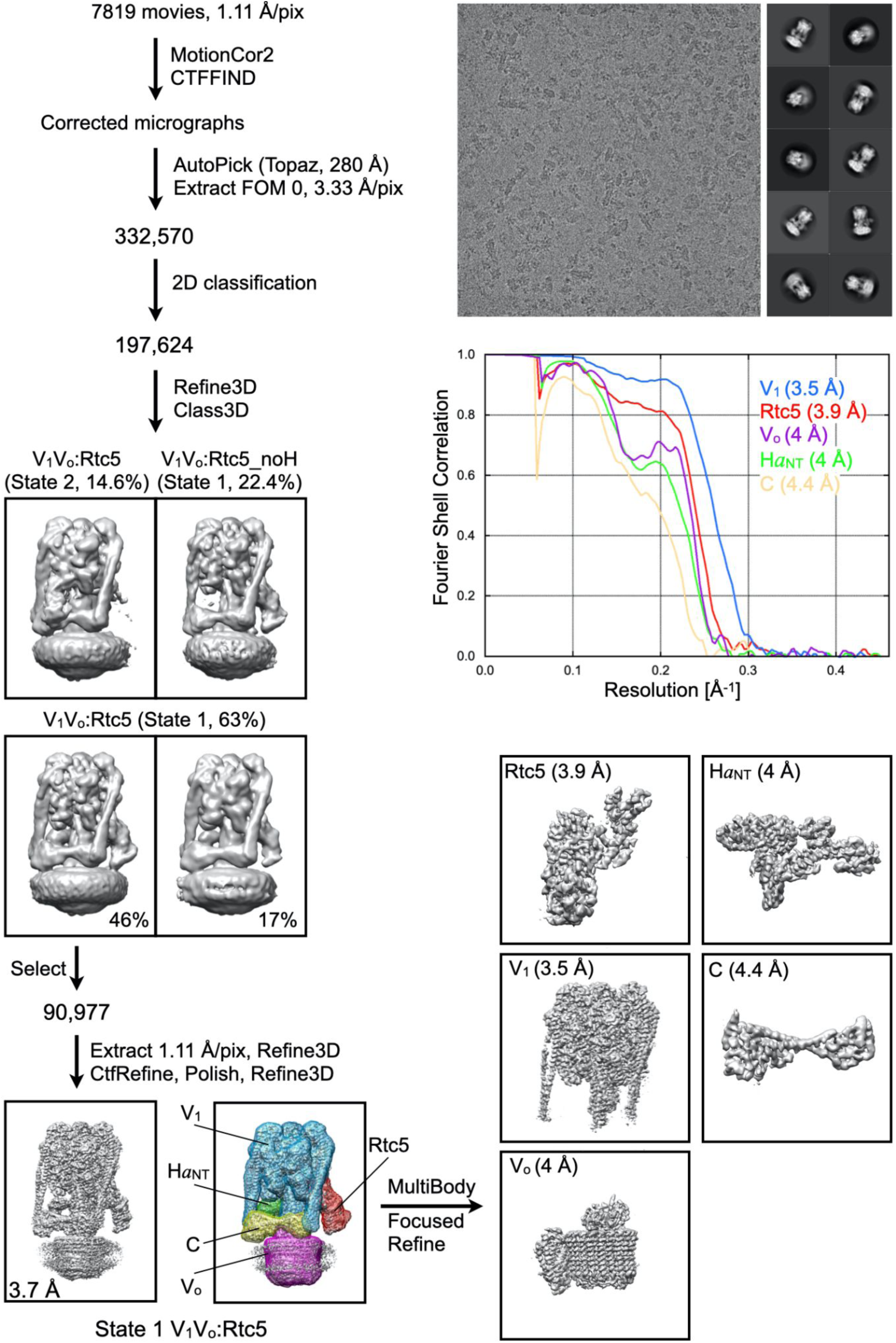
CryoEM structure analysis of Rtc5p bound post-assembly V-ATPase complex (V_1_V_o_ND:Rtc5^recon^). For details, see main text.

**Appendix Figure S4.**
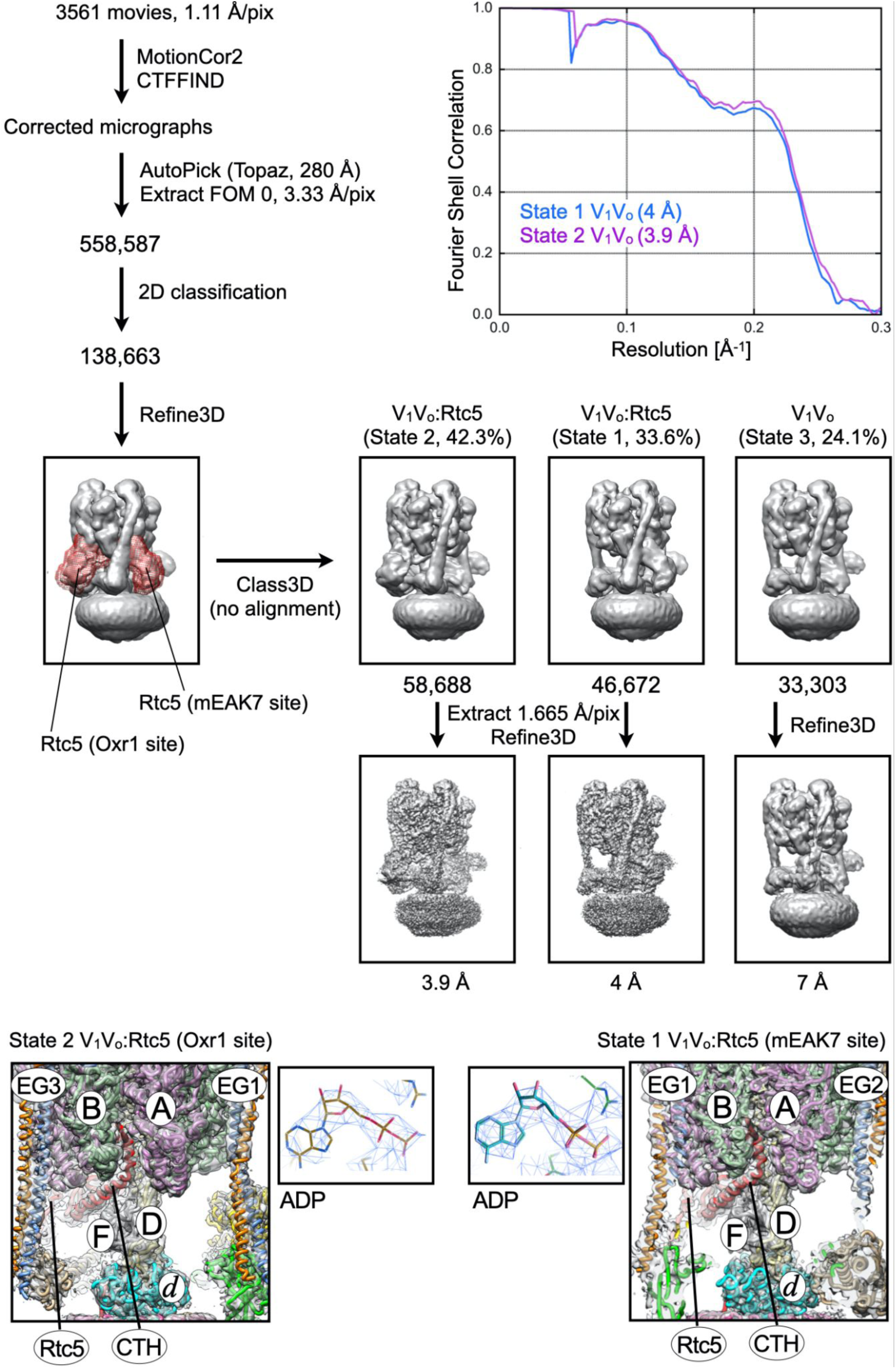
CryoEM structure analysis of endogenous V-ATPase vitrified in presence of added Rtc5p (V_1_V_o_ND:Rtc5p). For details, see main text.

**Appendix Figure S5.**
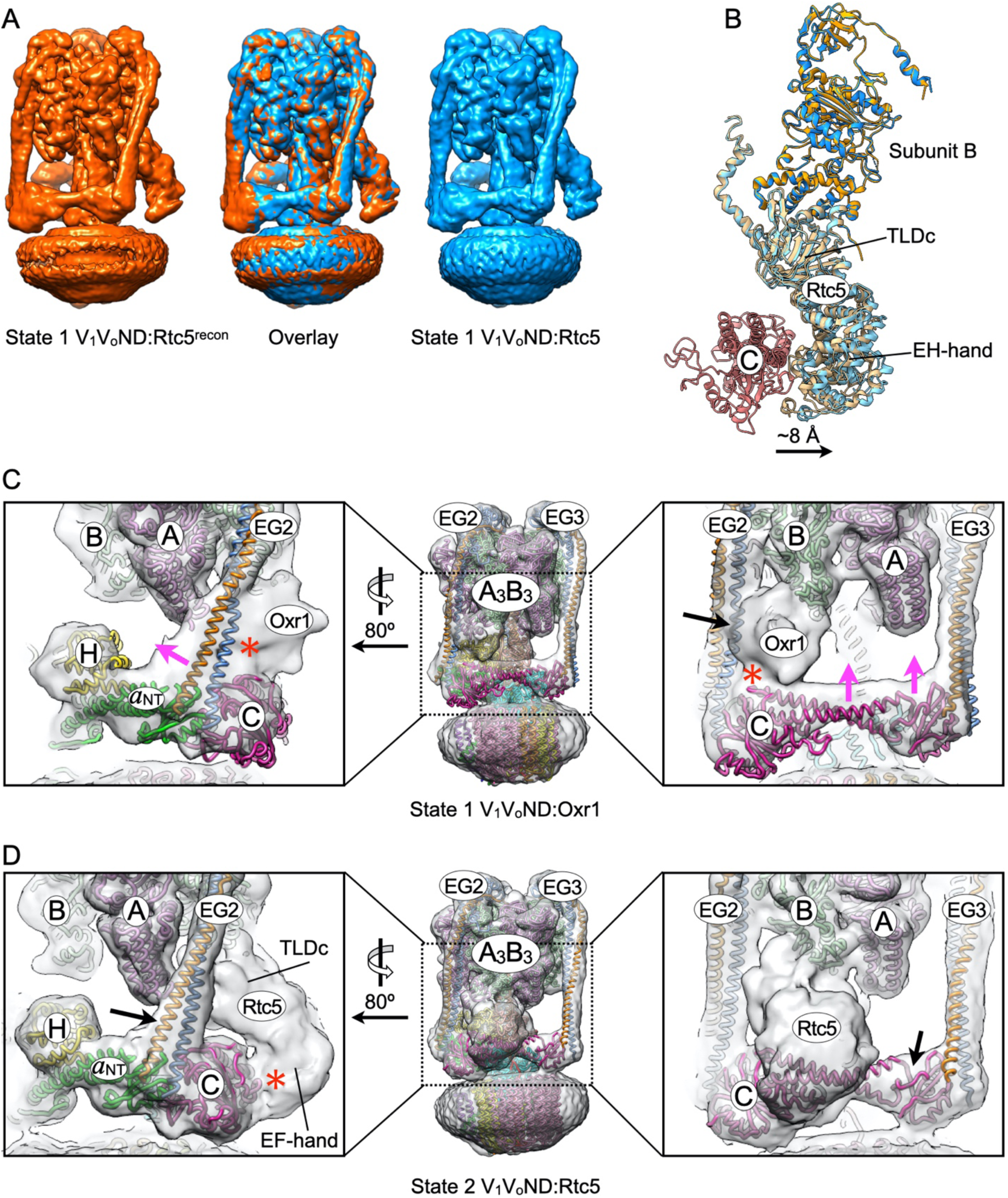
Comparison of state 1 V_1_V_o_ND:Rtc5^recon^ and V_1_V_o_ND:Rtc5, and impact of the interaction between state 1 V_1_V_o_ND and Oxr1p and Rtc5p. **A.** Overlay of state 1 V_1_V_o_ND:Rtc5recon (orange) and V_1_V_o_ND:Rtc5 (blue) maps. **B.** Comparison of the subunit B:Rtc5p sub-complexes of state 1 (blue) and state 2 (orange) V_1_V_o_ND:Rtc5. The sub-complexes were aligned using the B subunit coordinates (RMSD: 0.55 Å for the entire B subunit). **C.** Fitting of the coordinate file for state 1 V_1_V_o_ND (7FDA) into the map of Oxr1p bound state 1 V-ATPase (EMD-42880). Binding of Oxr1p leads to a ∼10 Å movement of subunit C towards Oxr1p (see pink arrows in the right panel) and a ∼20 Å movement of EG (see pink arrow in left panel). Oxr1p is seen to strongly interact with both EG2 (see red asterisk in left panel and black arrow in right panel) and the foot domain of the C subunit (see red asterisk in right panel). **D.** Fitting of the coordinate file for state 2 V_1_V_o_ND (7FDB) into the map of Rtc5p bound state 2 V-ATPase shows that binding of Rtc5p does not result in major structural rearrangements near its binding site (see black arrow in the right panel for subunit C and the left panel for EG2). In the state 2 V_1_V_o_ND:Rtc5 complex, Rtc5p’s EF-hand forms a large interface with subunit C (see red asterisk in the left panel).

**Appendix Figure S6.**
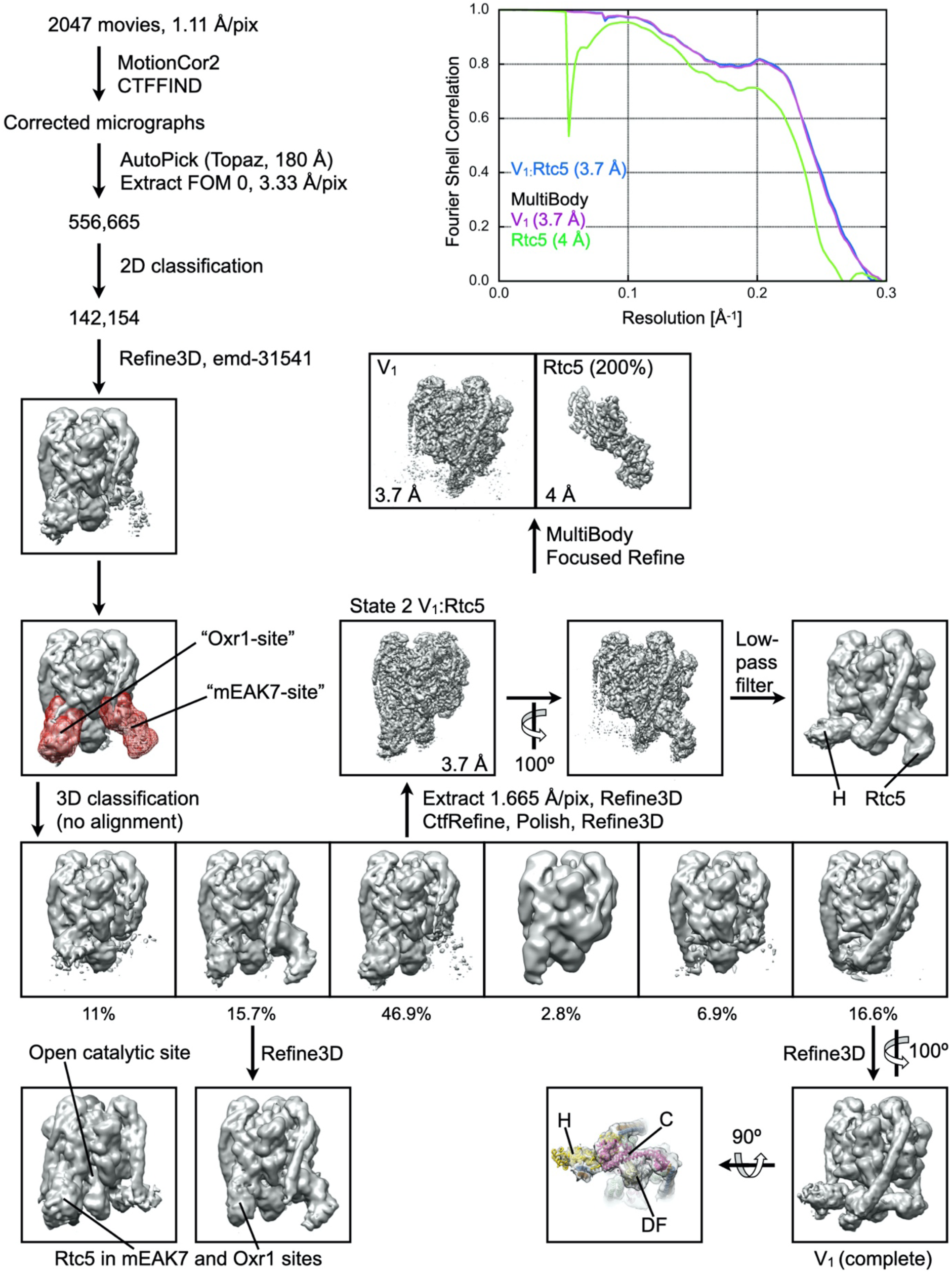
CryoEM structure analysis of the pre-assembly V_1_:Rtc5p complex. For details, see main text.

**Appendix Figure S7.**
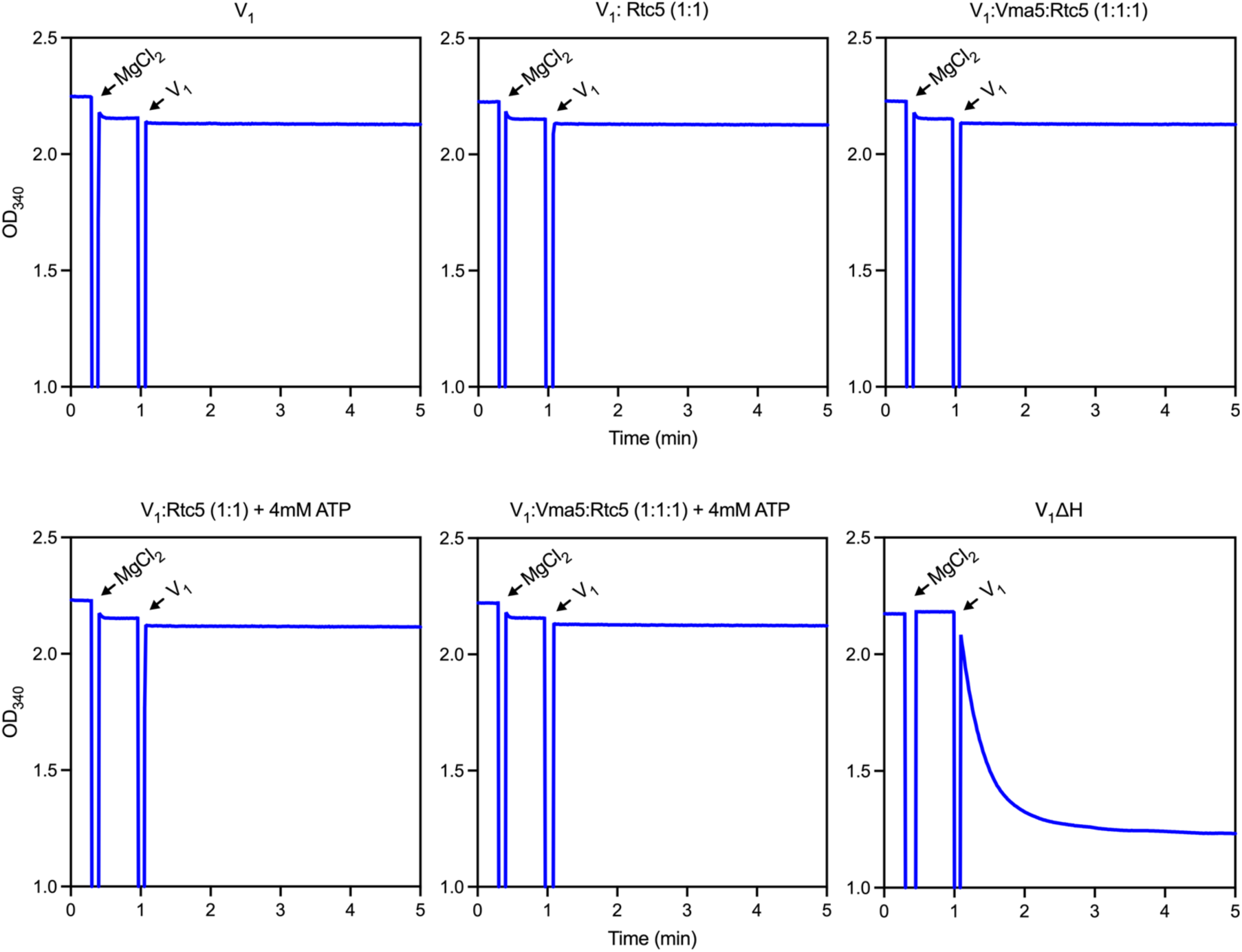
Rtc5p does not activate wild type autoinhibited V_1_ subcomplex. Purified wild type autoinhibited V_1_ subcomplex was incubated with either Rtc5p (top middle), Rtc5p and subunit C (Vma5p) (top right), Rtc5p and MgATP (bottom left), and Rtc5p, Vma5 and MgATP (bottom middle) in the indicated molar ratio or concentration for 2 h at room temperature before measuring the activity in an ATPase assay. As control, assay traces for autoinhibited wild type V_1_ (top left) and mutant active V_1_ (V_1_ΔH) (bottom right) are shown. Assays were supplemented with MgCl_2_ before adding V_1_ samples. A representative of six assays from two biological preparations is shown.

**Appendix Figure S8.**
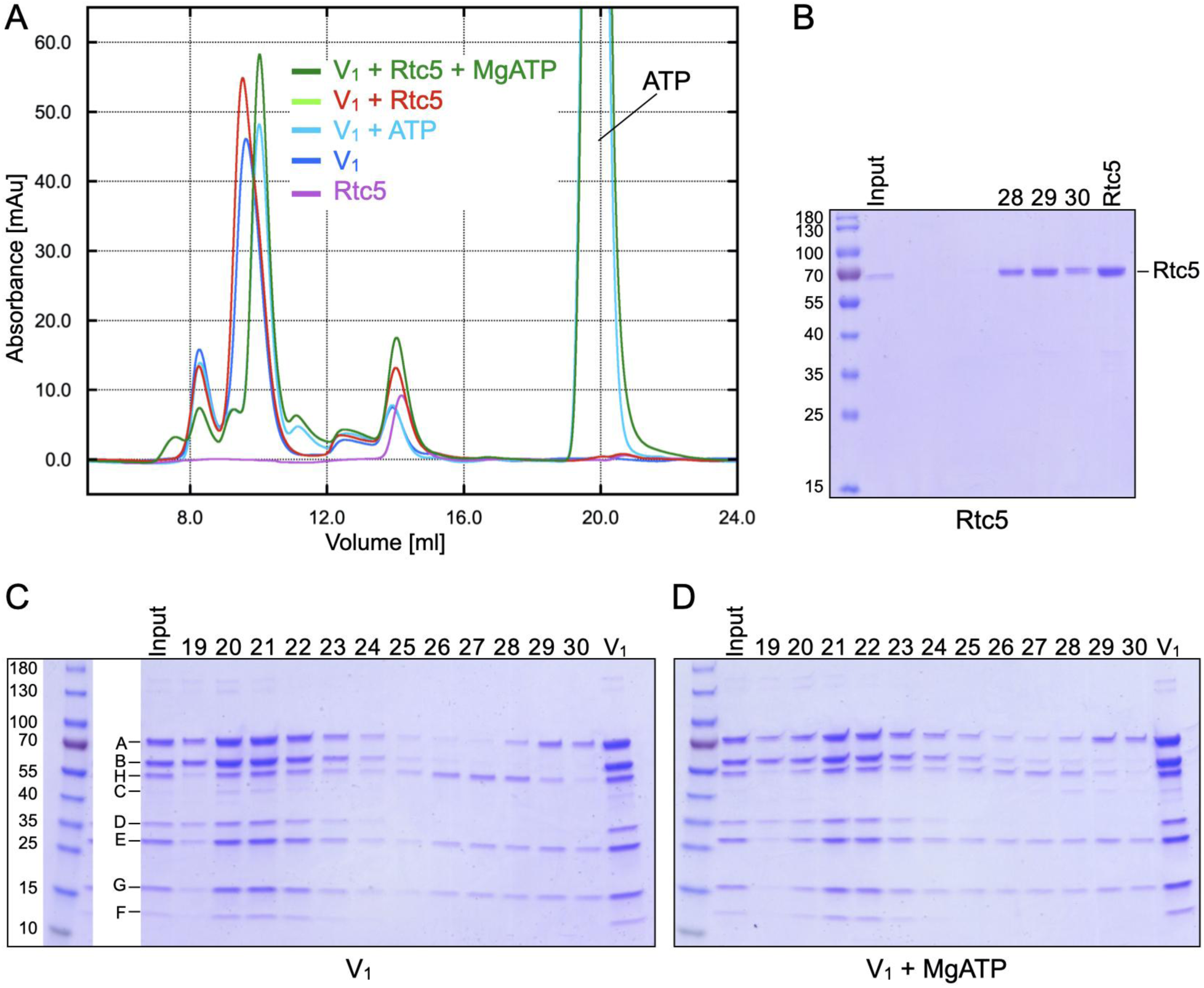
Size exclusion chromatography of Rtc5p and V_1_ in absence and presence of MgATP. **A.** Overlay of size exclusion chromatography elution profiles for V_1_ (blue), V_1_+MgATP (cyan), V_1_+Rtc5p (red), V_1_+Rtc5p+MgATP (green), and Rtc5p (purple). **B.** Coomassie Blue stained SDS-PAGE of Rtc5p elution fractions. **C.** Coomassie Blue stained SDS-PAGE of V_1_ elution fractions. **D.** Coomassie Blue stained SDS-PAGE of V_1_ + MgATP elution fractions. Representative gels of two biological repeats are shown. For the V_1_ + Rtc5p and V_1_ + Rtc5p SDS-PAGE gels, see main text.

**Appendix Figure S9.**
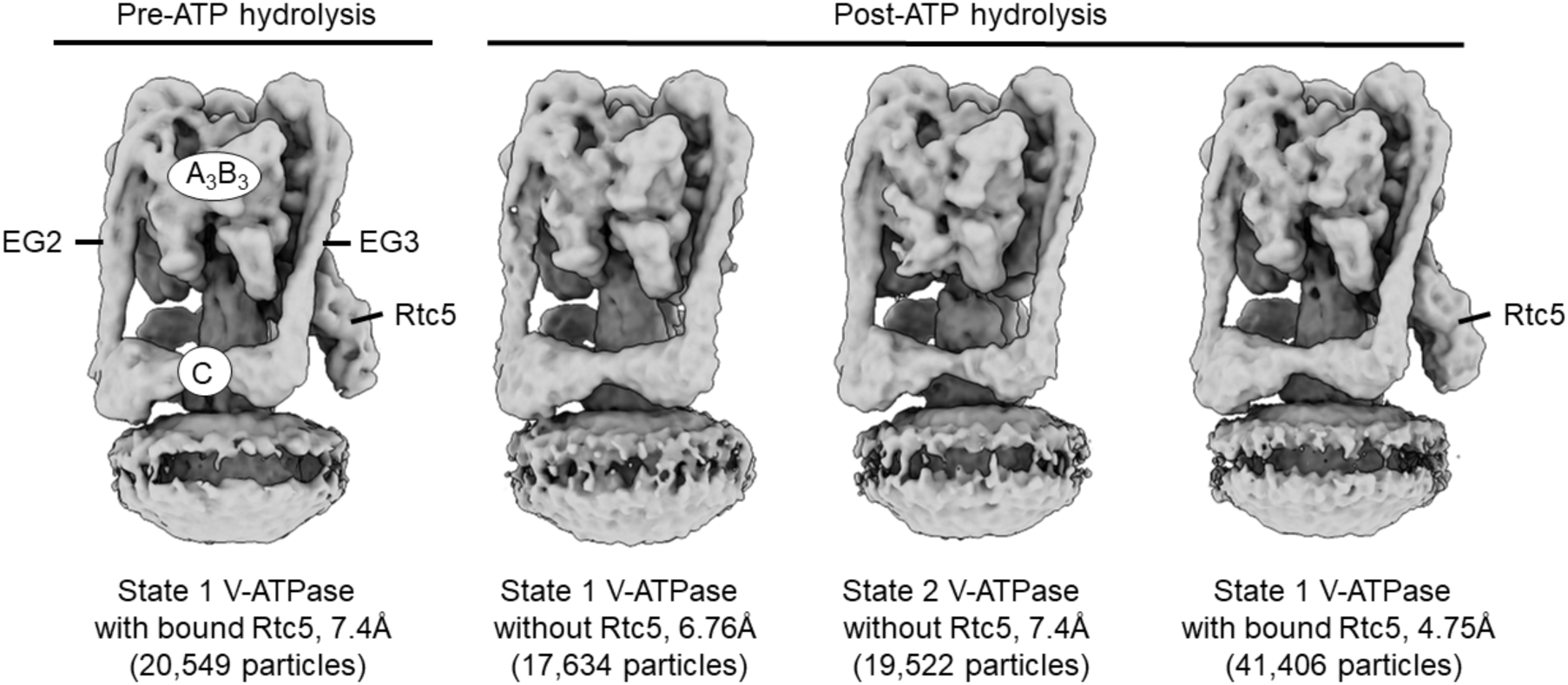
CryoEM maps of Rtc5p bound V-ATPase vitrified in presence of MgATP (V_1_V_o_ND:Rtc5^recon^ +/- MgATP) **Left panel** (pre-ATP hydrolysis), cryoEM map of *in vitro* assembled V-ATPase with bound Rtc5p in rotary state 1. **Right panels** (post-ATP hydrolysis), cryoEM maps of V_1_V_o_ND:Rtc5^recon^ vitrified in presence of 600 µM MgATP. ATP treatment results in three classes, with two classes free of Rtc5p (∼50% of the particles) in rotary states 1 and 2, and the third class with Rtc5p bound in rotary state 1.

**Appendix Figure S10.**
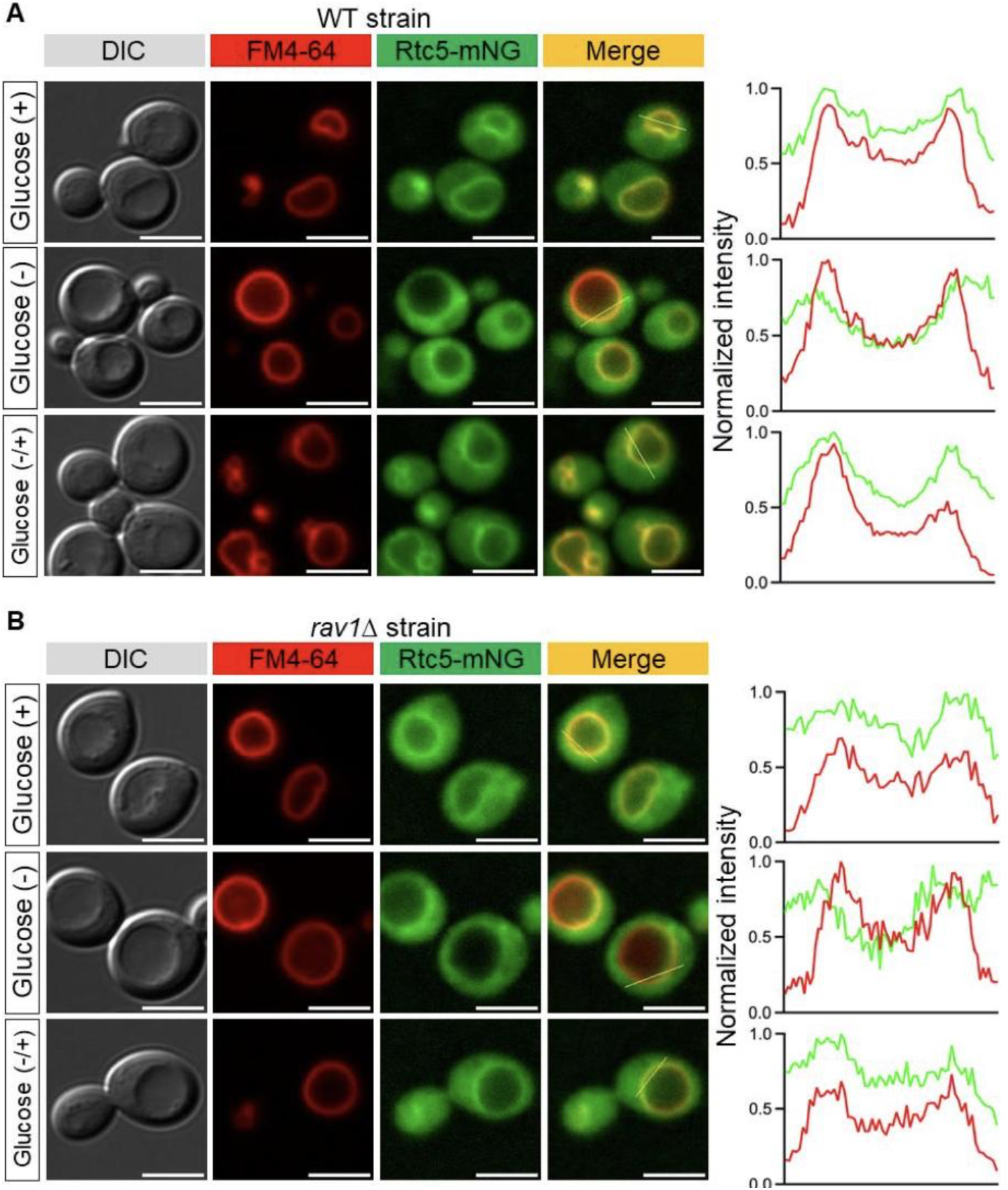
Localization of Rtc5-mNG in response to glucose. **A-B**. Yeast cells expressing C-terminally mNG tagged Rtc5p (Rtc5-mNG) in wild type (panel A) and *rav1*Δ (panel B) backgrounds were grown in glucose-containing media to exponential phase before depriving of glucose for 15 min, followed by 15 min of growth after re-adding glucose. Cells were withdrawn from each growth condition and immediately imaged by fluorescence microscopy. In both wild type and *rav1*Δ strains, Rtc5-mNG is on the vacuole in glucose-fed condition, becomes cytosolic in response to glucose deprivation, and returns to the vacuole after re-adding glucose. Vacuoles were stained with FM4-64 dye. Around 50-100 cells from two replicates were analyzed. Right panel shows the line scan profile of the merged image in each condition. Scale bar: 5 μm.

**Appendix Figure S11.**
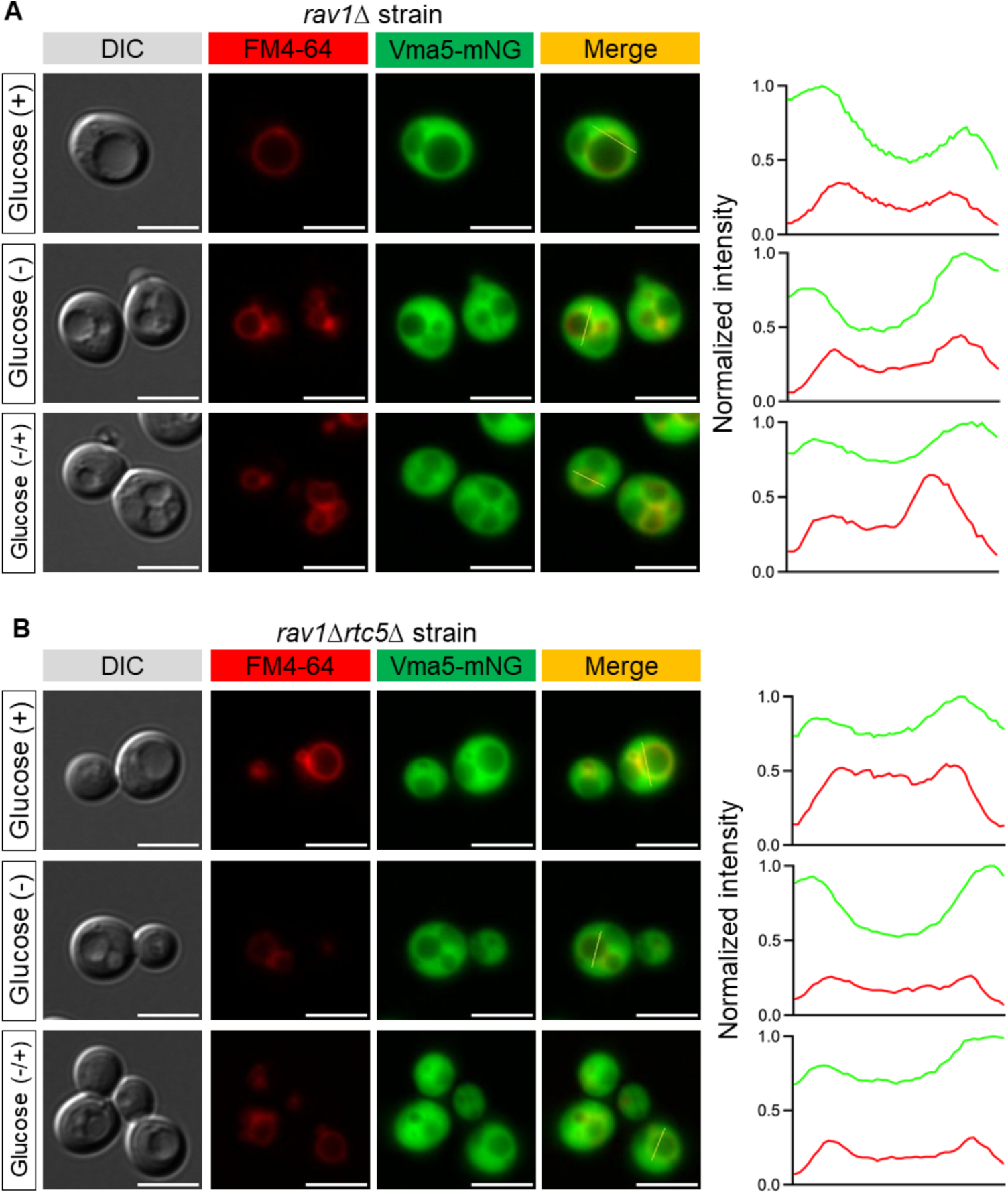
Reversible disassembly in *rav1*Δ and *rav1*Δ*rtc5*Δ strains. **A-B.** Yeast cells expressing C-terminally mNG tagged subunit C (Vma5-mNG) in *rav1*Δ (panel A) and *rav1*Δ*rtc5*Δ (panel B) backgrounds were grown in glucose-containing media to exponential phase before depriving of glucose for 15 min, followed by 15 min of growth after re-adding glucose. Cells were withdrawn from each growth condition, and immediately imaged by fluorescence microscopy. In both strains, Vma5-mNG is cytosolic in all growth conditions. Vacuoles were stained with FM4-64 dye. 50-100 cells from two replicates were analyzed. Right panel shows the line scan profile of the merged image in each condition. Scale bar: 5 µm.

**Appendix Table S1.** CryoEM data collection, refinement and validation statistics.

|  | V<sub>1</sub>V<sub>0</sub>ND:Rtc5<sup>recon</sup><br>(state 1) EMD-48491<br>PDBID: 9MOY | V<sub>1</sub>:Rtc5 (state 2)<br>EMD-49556<br>PDBID: 9NN1 | V<sub>1</sub>V<sub>0</sub>ND:Rtc5 (state 2)<br>EMD-70377<br>PDBID: 9ODU |
| --- | --- | --- | --- |
| Data collection and processing |  |  |  |
| Magnification | 92,000 | 92,000 | 92,000 |
| Voltage (kV) | 200 | 200 | 200 |
| Electron exposure (e<sup>-</sup>/Å<sup>2</sup>) | 52.5 | 49.25 | 49.25 |
| Defocus range (μm) | 0.4 - 2 | 0.4 - 2 | 0.4 - 2 |
| Pixel size (Å) | 1.11 | 1.11 | 1.11 |
| Movies recorded | 7819 | 2047 | 3561 |
| Symmetry imposed | C1 | C1 | C1 |
| Initial particle images (no.) | 332,570 | 556,665 | 558,587 |
| Final particle images (no.) | 90,977 | 61,771 | 58,688 |
| Map resolution (Å) | 3.7 | 3.7 | 3.9 |
| FSC threshold | 0.143 | 0.143 | 0.143 |
| Model building & refinement |  |  |  |
| Initial model used (PDB code) | 7fda, 7fdb, AlphaFold | 7tmm, 7fdc, 9moy | 7fdb, 9nn1, 9moy |
| Model resolution (Å) | 3.7 | 3.9 | 4.3 |
| FSC threshold | 0.5 | 0.5 | 0.5 |
| Model composition |  |  |  |
| Non-hydrogen atoms | 68,942 | 38,845 | 69,075 |
| Protein residues | 8,847 | 4,941 | 8,867 |
| Ligands | 1 | 1 | 1 |
| B factors (Å<sup>2</sup>) |  |  |  |
| Protein (min/max/average) | 55/329.2/119 | 56/1034/145 | 0/920/157 |
| Ligand | 86 | 120 | 111 |
| R.m.s. deviations |  |  |  |
| Bond lengths (Å) | 0.005 | 0.003 | 0.003 |
| Bond angles (°) | 0.638 | 0.484 | 0.487 |
| Validation |  |  |  |
| MolProbity score | 1.21 | 1.15 | 1.39 |
| Clashscore | 2.61 | 2.66 | 3.6 |
| Poor rotamers (%) | 0 | 0 | 0 |
| Ramachandran plot |  |  |  |
| Favored (%) | 97.07 | 97.53 | 96.44 |
| Allowed (%) | 2.93 | 2.47 | 3.56 |
| Disallowed (%) | 0 | 0 | 0 |
| Ramachandran Z-score (RMSD) |  |  |  |
| Whole | -1.28 (0.08) | -0.01 (0.12) | -0.93 (0.09) |
| Helix | -0.84 (0.07) | 0.46 (0.11) | -0.08 (0.08) |
| Sheet | 0.05 (0.2) | -0.35 (0.2) | -1.28 (0.19) |
| Loop | -0.79 (0.11) | 0.21 (0.14) | -1.07 (0.11) |

**Appendix Table S2.** Focused refinement of V_1_, V_o_, H*a*_NT_, Rtc5p and subunit C from state 1 V_1_V_o_ND:Rtc5^recon^ (post-assembly complex)

|  | V<sub>1</sub><br>EMD-48486 | V<sub>o</sub><br>EMD-48487 | Ha<sub>NT</sub><br>EMD-48488 | Rtc5<br>EMD-48489 | Subunit C<br>EMD-48490 | V<sub>1</sub>V<sub>o</sub>:Rtc5<br>Composite<br>EMD-48486 |
| --- | --- | --- | --- | --- | --- | --- |
| Data collection and processing |  |  |  |  |  |  |
| Magnification | 92,000 |  |  |  |  |  |
| Voltage (kV) | 200 |  |  |  |  |  |
| Electron exposure (e<sup>-</sup>/Å<sup>2</sup>) | 52.5 |  |  |  |  |  |
| Defocus range (μm) | 0.4 - 2 |  |  |  |  |  |
| Pixel size (Å) | 1.11 |  |  |  |  |  |
| Movies recorded | 7819 |  |  |  |  |  |
| Symmetry imposed | C1 |  |  |  |  |  |
| Initial particle images (no.) | 332,570 |  |  |  |  |  |
| Final particle images (no.) | 90,977 |  |  |  |  |  |
| Map resolution (Å) | 3.5 | 4 | 4 | 3.9 | 4.4 |  |
| FSC threshold | 0.143 | 0.143 | 0.143 | 0.143 | 0.143 |  |
| Map resolution range (Å) |  |  |  |  |  | 3.5 – 4.4 |

**Appendix Table S3.** Focused refinement of V_1_ and Rtc5p from state 2 V_1_:Rtc5 (pre-assembly complex)

|  | V<sub>1</sub><br>EMD-49557 | Rtc5<br>EMD-49558 | V<sub>1</sub>:Rtc5 (Composite)<br>EMD-49559 |
| --- | --- | --- | --- |
| Data collection and processing |  |  |  |
| Magnification | 92,000 |  |  |
| Voltage (kV) | 200 |  |  |
| Electron exposure (e<sup>-</sup>/Å<sup>2</sup>) | 49.25 |  |  |
| Defocus range (μm) | 0.4 - 2 |  |  |
| Pixel size (Å) | 1.11 |  |  |
| Movies recorded | 2047 |  |  |
| Symmetry imposed | C1 |  |  |
| Initial particle images (no.) | 556,665 |  |  |
| Final particle images (no.) | 61,771 |  |  |
| Map resolution (Å) | 3.7 | 4 |  |
| FSC threshold | 0.143 | 0.143 |  |
| Map resolution range (Å) |  |  | 3.7 – 4 |

**Appendix Table S4.**
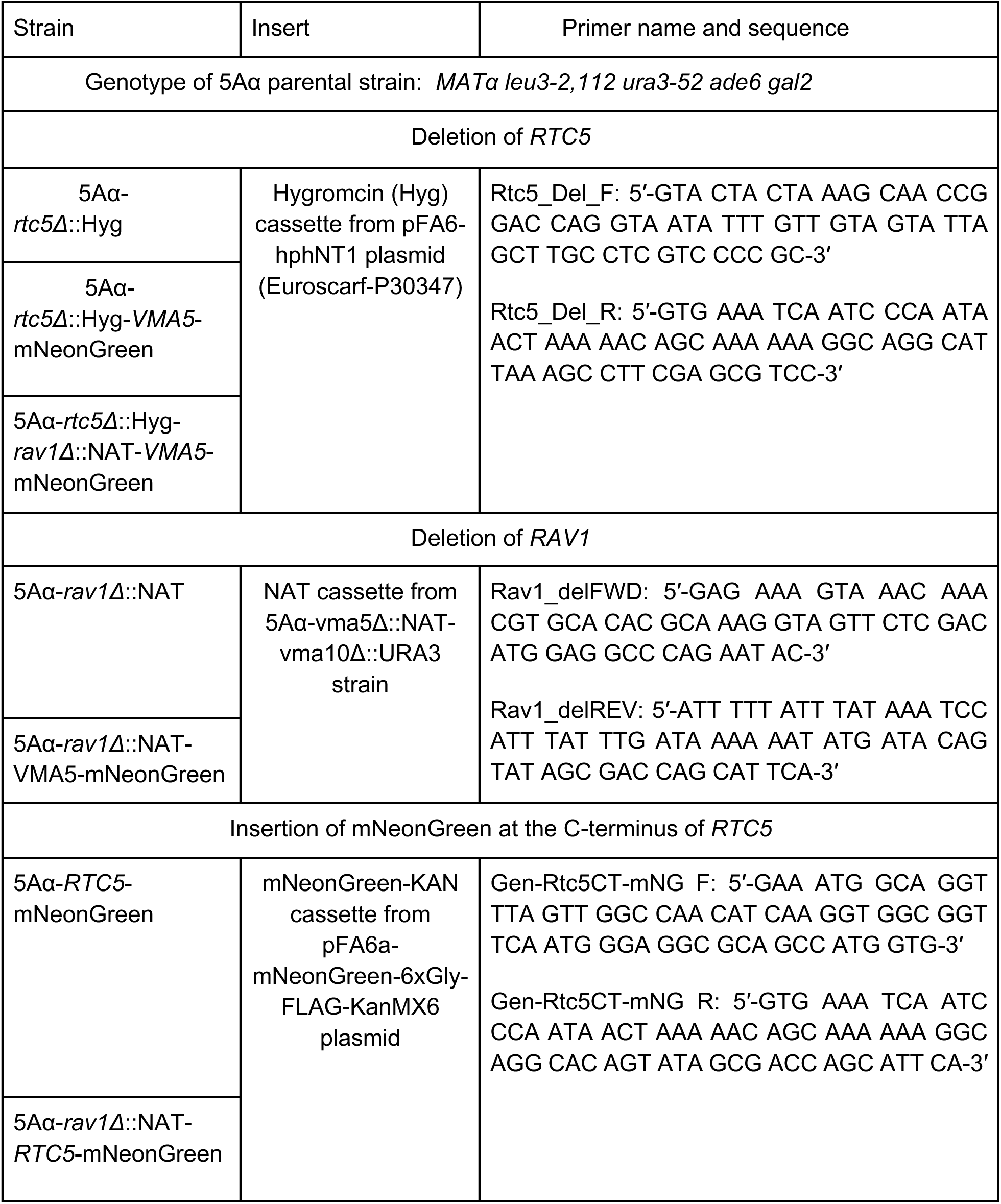
List of yeast strains generated in this study.

## Appendix Movies

**Appendix Movie S1**

**Appendix Movie S2**

